## Supplemental information for "Palaeogenomic analysis of black rat (*Rattus rattus*) reveals multiple European introductions associated with human economic history"

***Rattus rattus* supplementary for paper**

**Supplementary Notes 1: Site descriptions**

**Archaeological site descriptions**

We sampled 200 ancient black rat individuals from 34 archaeological sites across Europe, North and East Africa, and southern Asia (Supplementary Table 7). Where multiple samples were taken from the same or related archaeological contexts, care was taken to ensure that these represented discrete individuals - either by sampling the same skeletal element and side or on the basis of differing size and/or age.

***Althiburos, Tunisia*** *(Silvia Valenzuela-Lamas)*

The site of Althiburos (el Médéïna, Tunisia; longitude 8,790329, latitude 35,877177) is located in the present-day of Le Kef region, far from the sea (120 km to the S from Tabarka, 180 km to the SW from Carthage, 165 km to the W from Sousse), but well connected with the coast through the ancient road from Carthage to Theveste. It is a multi-period site with almost continuous occupation from the 9th century BCE to 13th century CE [1]. Many artifacts were recovered during the excavation fieldworks, including a vast faunal assemblage with diverse terrestrial fauna spanning the whole diachrony of the site [2]. The area 1, where the rat specimens come from, is located adjoining the Roman capitolium of the city and comprised several rooms, probably of domestic use [3]. More precisely, SU 190016, from which the rat bones were recovered, corresponds to the grey filling of a gutter built in Roman times. The pottery associated with the rat bones – Late Roman C ware and an edge of regional sigillata form sim. Stern IV – suggested a probable Late Roman chronology, but radiocarbon dating of one of the sequenced specimens (ATU001) gave a calibrated range of 706-883 CE (95.4%), thus suggesting that our samples date from the 8th or 9th century CE.

***Castle of Aqaba, Jordan*** *(Wim Van Neer)*

The town of Aqaba, located on the Red Sea coast in southern Jordan, counts several historical sites that were investigated archaeologically. One of them is the castle that was probably built in the 12th century CE by the crusaders. Excavations were carried out there between 2000 and 2008 by the Ministère de la Région wallonne (Belgium), Cardiff University (UK) and Ghent University (Belgium). The recovered faunal remains date between the Islamic and Modern period, with most of the finds dating to the Mamluk and Ottoman periods [4]. The sole black rat find from the site is a complete skeleton dated to the Mamluk period (late 13th – 15th century CE) from which a femur was taken for aDNA analysis.

***Assos, Turkey*** *(Beate Böhlendorf-Arslan)*

Assos, located in the southwest of the Anatolian peninsula and opposite the island Lesbos, was founded by Greek colonists from Methymna in the 7th century BCE [5]. Starting from the second half of the 5th century, the antique city was remodeled [6,7]. During the transformation of the city, a terrace south of the agora, the old city center, was converted and provided with a church, a reception hall and other buildings.

The teeth of two rats were an unexpected find and came from the flotation material of two pithoi. This material, a tangle of archaeobotanical remains, was transferred to Germany for analysis with an official permit in 2015. It was taken from a building nearby the reception hall that may have been part of a bishop's palace [7,8]. Two of the rooms in this L-shaped building served as depots for building materials and for storing food, which were placed in the two large pithoi. Among the finds in this room, in addition to pottery and small finds, were coins, including a bronze half-follis of the emperor Leontius (695-698 CE) [9]. This coin gives a clue to the destruction of the building, whose roof collapsed at the very end of the 7th or beginning of the 8th century CE, burying the inventory.

Both teeth were sampled, and both had high enough endogenous DNA content to be included in the analysis (Ass_1, Ass_2).

***Buda Castle - Teleki Palace, Hungary*** *(Zsófia E. Kovács)*

Buda Castle was the medieval royal capital of Hungary (situated on the right bank of the Danube). The Ottoman occupation of the city was in 1541 and lasted until 1686. The city of Buda was an important administrative and commercial centre of the Ottoman Empire.

Excavation of the remains of the so-called Teleki Palace in the south part of the Buda Castle district, were directed in 1998-2000 by Dorottya B. Nyékhelyi [10,11]. Animal bone materials came from numerous pits and a well dated to the medieval and Ottoman period based on archaeological stratigraphy and associated archaeological finds [10–12]. Many black rat finds were recovered from the hand-collected material of the site [13]. Rat bones (three tibias and three femurs) analysed in this study derived from medieval levels (14th-15th century CE) of three pits (pit 11, 15, 16), and from a pit (pit C5/7) dated to the Ottoman period (16th-17th century CE).

***Caričin grad, Serbia*** *(Henriette Baron)*

The fortified settlement of Caričin grad (near Lebane in southern Serbia) is confidently associated with this historical known city of Justiniana Prima, founded by Justinian in 535 CE and abandoned around 615 CE. Recent excavations by the Archaeological Institute in Belgrade and the Römisch-Germanisches Zentralmuseum in Mainz included an intensive programme of flotation for botanical and small vertebrate remains. A large quantity of rodent bones, dominated by black rat, was recovered from deposits associated with a large granary[14].

A total of 32 specimens were screened for aDNA. All but four of these were selected for deeper sequencing, with four ultimately included in the nuclear genomic study (Car_4, Car_5, Car_7, KR150770) and 28 contributing mitochondrial data.

***Chersonesos, Ukraine*** *(Pavel Gol'din)*

The city of Chersonesus (also known as Chersonesos Taurica or Tauric Chersonesos) (44°36'39"N, 33°29'39"E) was the largest city and the greatest trade and cultural centre of the northern Black Sea region during the Hellenistic, Roman and Byzantine time. Founded as a Greek colony in the 5th century BCE, it was historically in the orbit of Pontus, Bosporus and Rome. The Northern District of the city was among the earliest parts of the city housing the port area, fish market, fish salting facilities and temples [15,16]. Block 9A, where the rat specimens come from, was the location for the temple complex, city square, living houses and numerous wells [16]. The temple complex possibly functioned between the 4 century BCE and 4 century CE. Many artifacts, mostly of the Roman and early Medieval time, were excavated from this area in 2012-13 [16]), as well as kitchen waste containing faunal assemblage with diverse marine and terrestrial fauna.

Four rat specimens were screened, deriving from 1m-4m depths within a well excavated in 2005. They were dated to between the 1st C BCE and the 1st century CE based on archaeological stratigraphy and associated finds. However, a radiocarbon date on a fifth rat bone from c.4m depth in the same well gave a calibrated range of 130-320 CE (95.4%), suggesting that our samples are all from the 2nd century CE or later.

***Chombo, Kenya (Richard Helm)***

CHO 01 TP04 (13): Chombo Site 01 Test Pit 04 Context 13

CHO 01 TP04 (14): Chombo Site 01 Test Pit 04 Context 14

Chombo is an open settlement site, situated at the base of the Shimba Hills, Kwale County in coastal Kenya. The site occupies a tributary valley of the Cha Simba (Pemba) river. The two rat specimens derive from a single test pit excavated in 1997 [17]. The site represents a rural farmstead potentially occupied from as early as the eighth century and continuing until at least the late tenth century. The specimens were recovered from a consecutive sequence of habitation surfaces (ash, charred plant remains, animal bone and shell), and included evidence for iron working residues (slag and tuyere fragments). A charred wood sample from the layer immediately below the earliest recovered rat specimen returned a radiocarbon date of 1180 ± 60 BP (cal AD 867-986; Pta-7978).

***Deventer - Stadhuiskwartier, Netherlands*** *(Inge van der Jagt)*

Deventer is one of the oldest cities in the Netherlands. The city has a port and is located on the river IJssel in the middle of the Netherlands. Due to its strategic location, it played an important role in medieval trade, including the Hanseatic trade from the end of the 13th centuries onwards.

The remains of the rats were found at the site Stadhuiskwartier (this can be translated as city hall quarter). The finds of the excavations at this location provide a diachronic picture of the settlement history of Deventer. The oldest phases date from the Mesolithic and the Late Neolithic. After that, there are occupation layers from the Bronze age, Iron age and Roman period. From the 8th century onwards the habitation is uninterrupted and gradually changed from an agricultural to a more urban character. During the 9th century trade and craft production became increasingly important. After 882 AD an earthen defense wall was constructed in response to a Viking attack, which is seen as an important next step in the urbanisation of Deventer. From the 11th century the Stadhuiskwartier was the residence of the powerful of the city. It is therefore not surprising that from this time onwards this area of the city also became the center of urban administration.

The samples are taken from three different cess pits. They are all square, brick-lined cesspits. The oldest cesspit (cesspit 19) dates back to 1350-1400 AD. It is likely that the pit belonged to the town wine house, the "Steerne". The last phase of use of this pit dates back to the second half of the 14th century. It contained four fragments of *Rattus rattus* (black rat) (sample 296). More or less simultaneously in use is cesspit 31 dating from 1350-1425 AD. The pit dates from just after the moment the plot came into the possession of the city, and became part of the city hall complex. The find material indicates that the owners were reasonably wealthy. In this cesspit the remains of 17 rats bones (*Rattus* sp.) were found (sample 758). The third cesspit (cesspit 18) has a much younger date (1620-1650 AD) and is located on the same plot as cesspit 31, which was no longer in use at that time. The pit was probably built in the 15th or 16th century, the last phase of use (from which the bone material also comes) dates from the second quarter of the 17th century. The cesspit cannot be attributed to specific residents or users, but the pottery and glass in the filling indicate an above-average prosperity. In the cesspit 22 remains of *Rattus* sp. were found (sample 745). All three cesspits contain many other animal remains in addition to the rat remains. This concerns not only domesticated species but also many remains of small wild mammals such as hedgehogs and Leporidae and many wild birds, including birds of prey such as a buzzard and kestrel in cesspit 19, and fish.

A total of 8 specimens from this site were screened for aDNA, of which 3 femurs were selected for deeper screening: one from cesspit 19 (SNE001), one from cesspit 31 (SNE002) and one from cesspit 18 (SNE004). The sample from cesspit 18 was directly radiocarbon dated to AD 1527-1794 at 95.4%

probability (MAMS-46370, 263±19), corroborating the archaeological dating. The dating of another sample from cesspit 31 failed because of insufficient collagen (Wl-51521).

***ed-Dur, United Arab Emirates*** *(Wim Van Neer)*

The site of ed-Dur is located on a plateau along the lagoon of Umm al-Qaiwain (UAE). It was inhabited mainly during the first and second centuries of the first millennium AD. Human and some animal burials have been found as well as a rather small number of building remains. Ephemeral habitation structures were probably present as well, judging from the abundant animal bones, pottery, glass and other objects found all over the site's surface. The presence of several objects of foreign origin and the favourable location of the site suggest that ed-Dur functioned as a harbour in international sea trade.

Faunal remains were collected during excavations carried out between 1987 and 1994 by an international consortium of teams from the universities of Ghent, Copenhagen, Lyon, Edinburgh and London [18]. Among the rodent remains, 23 rat bones were identified of which two femurs were selected for aDNA analysis. They belong to the first occupation phase of the site, dating between 50/25BC and AD150.

***Fort Frederik Hendrik, Mauritius*** *(Wim Van Neer)*

The island of Mauritius was known to Arab traders and to the Portuguese from at least the 13th-16th centuries, but the Dutch were the first to settle here in the 17th century. They built a fort that was excavated between 1997 and 2005 within the framework of the Frederik Hendrik Archaeological Project initiated by the Amsterdam Archaeological Center and the State University of New Jersey in collaboration with the Mauritius Institute, the National Heritage Trust of Mauritius and the Mauritius Museums Council.

A total of 74 black rat remains representing at least 8 individuals were found in layers dated to the last quarter of the 17th century [19]. Two rat femurs were sampled for analysis. According to historical records, the rat populations had devastating effects on the crops the Dutch tried to grow. Black rats were mentioned the first time in Dutch chronicles in 1606 but must have been present on the island before the first contact in 1598 as shown by subfossil remains and rat-predated snails. A Portuguese shipwreck was mentioned by the first Dutch visitors which may have been the source of these rats.

***Gatehampton Roman villa, England, UK*** *(Thomas Walker)*

The villa at Gatehampton is situated on a gravel terrace on the north side of the River Thames in south Oxfordshire. The villa forms part of a rural Roman farmstead, which was established in the second century and was under continuous occupation into the late fourth and perhaps early fifth century. It was excavated between 1993 and 2019 by the South Oxfordshire Archaeological Group (SOAG). The six rat femurs sampled here were part of a large assemblage of small mammal and other small vertebrate bones excavated within a 10 $m^2$ area of collapse debris in one room in the western end of the villa. The assemblage of bones, representing a minimum of 935 individuals including 14 individual black rats, is likely to derive from the pellets of barn owls which roosted in the building for one or two years between abandonment and collapse. Radiocarbon dating of additional rat specimens from the same deposit returned dates of 1760 ± 30 BP (cal AD 230-385; Beta-37717) and 1755 ± 27 BP (cal AD 235-380; UBA-40928). Contextual evidence for the abandonment of this section of the villa suggests that they accumulated during the middle or later half of the fourth century [20].

***Kalba, United Arab Emirates*** *(Alison Crowther)*

Kalba is a Bronze Age (Hafit or Umm an-Nar/Umm al-När period) settlement on the east coast of Sharjar in the Gulf of Oman. It was occupied from the third to mid-first millennium BCE. The site complex comprises a main settlement mound (Kalba Site 4, or K4), measuring ~50m in diameter and ~2.5m high, and several stone cairn tombs (K1A, K1B and K2) spread over an area of c. 40km2 surrounded by cultivated fields, coastal mudflats, and mangroves. Excavation of the mound by the Kalba Archaeological Research Project [21] revealed that occupation of the site was focused on a central, sub-circular mud-brick tower. There is also evidence of date palm production, animal husbandry (mostly caprines although cattle and equids were also present) and the use of marine resources.

A total of 47 rat bones were recovered from 17 contexts at Kalba [22] . These were concentrated in early second millennium BCE deposits and mostly associated with middens, as well as a significant proportion of rodent-gnawed bone refuse. Two bones were used for ancient DNA analyses, from contexts K2a 0007.9 and K2a 0007 upper 65m bay 32.

***Kantharodai, Sri Lanka*** *(Wijerathne Bohingamuwa)*

Kantharodai, located approximately 8 km from the ancient seaport Jambukolapattahana on the northern coast of Sri Lanka, was one of the four main urban cum religious centres of the island during the Early Historic period. It was invariably connected with the capital Anuradhapura, as well as ports across the Bay of Bengal. At Kantharodai, two trenches were excavated by the Department of Archaeology, Sri Lanka and floatation sampling was carried out by the Oxford University-based Sealinks Project in 2011 [23]. Rat bones were recovered from these flots. The combined site sequence here dates from c.400 BCE to 50/70 BCE. The rat bone sample from this site comes from Phase IV

(57-353 BCE), which has a median date of 178 BCE. The site appears to have declined just before the commencement of the Common Era.

***Karksi Castle, Estonia*** *(Eve Rannamäe)*

Karksi Castle is situated in the area of historical Livonia, present-day south Estonia. It was constructed on a formerly uninhabited site in the first quarter of the thirteenth century, as a result of the crusades [24]. The castle lost its importance during the Livonian War (1558–1583) and the Swedish-Polish wars (1600–1625). Excavations at Karksi Castle were carried out in 2011–2012 within the framework of the international project *The Ecology of Crusading* (European Union's Seventh Framework Programme FP7/2007–2013 under grant agreement no 263735). The excavation results are published in several articles and book chapters [24–28].

Faunal remains recovered during the excavations are abundant (over 11,500 specimens analysed [29]). Among the faunal assemblage from Trench 1, Layer 12, remains of common domestic animals, a few wild mammals, birds, various fish, and 27 rodent specimens were recorded. Two of the rodent bones – right *Rattus* femurs from Squares C/4 (specimen ID: TÜ-1929/2012/AZ-1:233) and C/7 (TÜ-1929/2012/AZ-7:2132) – were screened for DNA content (Supplementary Table 7); sampling protocol TÜ PP No 89, zooarchaeological collections of the Department of Archaeology, University of Tartu, Estonia.

The bones come from a partly waterlogged layer of black highly organic sediment that was very rich in both artefacts and ecofacts. The layer was interpreted as a midden (rubbish) deposit of a household and table waste, formed sometime during the last third of the 13th century, between 1266–1290 cal CE [25,30]. In Estonia, this was the beginning of the Middle Ages (c. 1225–1560) – the time of new power structures and construction of the castles, development of towns, introduction of Christianity, and changes in agriculture [27]. Taking into account the density of the midden, the very short timeframe given by the radiocarbon dating, and that the midden was rapidly buried [30], we could assume that the rat bones under question are indeed from a secure context of the last third of the 13th century.

***Kastelholm, Åland, Finland*** *(Hanna Kivikero)*

The castle of Kastelholm is located on the Åland Islands in the northern part of the Baltic Sea. The castle is first mentioned in 1388 and is thought to be founded between 1384 and 1387 [31,32]. The castle functioned as an administrative centre for the Swedish Crown until the early 17th century which started a slow process of decay of the castle [33]. After a fire in 1745 the castle was used solely for the storage of cereal and was no longer permanently occupied [34].

The analysed samples in Kastelholm come from three areas that were excavated between 1983 and 1985. One of the areas (KS5) was outside the north-eastern wall and the other two (KS8 and KS35) were located in the castle courtyard. The rat bones were primarily hand collected or sieved with 10 mm mesh.

A total of six rat specimens were submitted for aDNA analysis, two from each of the three areas. Two specimens selected for deeper sequencing derived from KS5 and KS8 (dated to c.1400 CE and 1400-1500 CE respectively based upon stratigraphy). Both of these specimens were then directly radiocarbon dated, and returned date ranges of 1422-1456 CE and 1328-1420 CE respectively at 95.4% probability (WK-51519, 463±21; WK-51518, 569±21).

***Kilton Castle, England, UK*** *(Terry O'Connor)*

Kilton Castle is in the North-East corner of Yorkshire (NGR NZ701175). Built in the 12th century on a strongly-defended promontory, it changed hands between local noble families and was ruinous by the 14th century. The castle was seized by the Crown following the Pilgrimage of Grace (1536-37) and was described as totally abandoned by the late 16th century. Excavations in the 1960s included a well, from the lower levels of which a mixed assemblage of rodent and felid bones was recovered [35], apparently by sieving. Deposits were wet and apparently anoxic: bone preservation is excellent. Bones from the well were taken in hand by Jennie Coy, who passed them on to Terry O'Connor in the early 1980s, describing the context as "from the Civil War" (i.e. 1642-1651). Subsequent correspondence with the excavator, the late Arthur ApSimon, indicated that the bone material was available for further research but did not further clarify the date. Given the known history of the building a Civil War date is unlikely and the filling of the well is more likely to derive from episodes of ruination in the 14th century. This is supported by a direct radiocarbon date of 1303-1402 CE at 95.4% (OxA-36673, 604±23) on a left rat femur.

A total of 20 right rat femurs were screened for aDNA, but only one was selected for deeper sequencing: KLT001.

***Mantai, Sri Lanka*** *(Wijerathne Bohingamuwa)*

Mantai (ancient *Mahatittha*/ *Mantottam*) is located on the northwestern coast of Sri Lanka. It was the main sea port of the capital, Anuradhapura, during the Early to Middle Historic periods of the island and was a vital sea link of the Indian Ocean maritime trade [36]. Mantai has an uninterrupted cultural sequence from *c.* 200BCE and up to 850CE, after which the site is highly disturbed. These upper levels, however, contain quantities of imported ceramics that are clearly datable up to the 12th/13th centuries, indicating the continuity of the port [23,37].

The rat bones were sampled during the Oxford University-based Sealinks project excavation in 2009/2010. They come from Phase I {(3326+/-38 BP) *c.* 1600 BCE} and IV {(1510=/-37BP) 460-610 CE; by=250-380CE)}. The earliest phase here can tentatively be assigned to late Mesolithic period, when the site was briefly occupied and abandoned. Phase IV is the Middle Historic period during which Mantai started to gain prominence as an International seaport cum port city [23,37].

***Mértola, Portugal*** *(Arturo Morales)*

The site of Mértola (Baixo Alentejo, Portugal) is located on the top of a hill on the eponymous city that borders the Guadiana river. During the Islamic period (8-13th C CE) the city was surrounded by a defensive wall. The rat remains were retrieved in one of the tightly packed houses that constituted the living quarters at that time [38].

Contextually speaking, the remains derive from two units of this house, (a) the Telhao unit, a living floor, sealed under a collapsed roof, and (b) the Fosa unit, a 1m wide X 1.5m deep oval cesspit lying adjacent to the external wall of this house on a 2m wide street. The cesspit was sealed by a tightly fitting slab and was a waste disposal area. The materials were dated to the first quarter of the 13th C AD based upon archaeological stratigraphy and associated finds. This is just prior to the Christian occupation of this stronghold (from 1328 CE), coincident with the Almohad occupation of southern Iberia. After the Christian occupation, Muslim cesspits were abandoned and many lost the slabs that covered them, which then allowed intrusive materials to mix with the original deposits. This does not appear to be the case for the Fosa unit, although the possibility that some or most of these rats represent later intrusives cannot be excluded as the cesspits could be reached via subterranean galleries.

A total of 11 specimens were selected for aDNA analysis, two from the Telhao and nine from the Fosa. All were right tibiae. Of three specimens selected for deeper sequencing, one sample from each context was also directly radiocarbon dated to AD 1161-1259 and AD 1166-1261 respectively at 95.4% probability (Wk-51525, 852±21; WK-51526, 843±21), corroborating the archaeological dating.

***Castle of Nürnberg, Bavaria, Germany*** *(Kerstin Pasda)*

The castle consisted of two different areas: The Palatinate and the residence of the magistrate of the castle (“Burgamtmannsgebäude”). The dating of the bone material reached from the 11$^{th}$ to the 15$^{th}$ century. The castle of Nürnberg ranks among the most significant castles in medieval central Europe. Frequent stays of emperors and kings until 1571 are recorded and hence demonstrate, that the castle of Nürnberg was a place of eminent political events. Since the mid of the 11$^{th}$ century, a castle which was in the possession of Salian kings existed in Nürnberg [39]. The emperors Heinrich III (1039-1056), Heinrich IV and Heinrich V were owners of the castle. Nürnberg became imperial place in 1138 under the Staufer king Konrad III (1138-1152). This was also the first time, when a spatial and juridical division of Palatinate and residence of the magistrate became visible. Again, a change of ownership took place in 1192, when the castle went to the earls of Hohenzollern. The famous Friedrich II who wrote the earliest book about falconry ("de arte venandi cum avibus") stayed at the castle between 1210-1220. Under the king Rudolf I of Habsburg (1273-1291) the castle exhibited again significance. The influence of the town Nürnberg increased under the authority of emperor Heinrich VII in 1313 and the whole castle came into possession of the town in 1427.

Nobody stayed permanently at the Palatinate. Living rooms were used during the stay of the current monarch. The hall of the Palatinate was used for diets and congregations. The residence of the magistrate was occupied permanently at the latest from 1138 on when the counts von Raabs from Lower Austria became the first magistrates [40,41].

The bone material of the Palatinate was hand collected, the material of the residence was sieved mainly. All together 18829 animal bones and 3 human bones were analysed in a thesis [42]. In the excavation of the Palatinate 3612 bones and in the residence 13230 bones were analysed. All together 25 bones of rats were identified. One bone sample of a rat, which was found in a layer of the residence and which was dated to the second half of the 12th century was used in the project.

***Monte di Tuda, Corsica, France*** *(Jean-Denis Vigne)*

The small Monte di Tuda cave (Olmetta-di-Tuda) is located in the North of Corsica (France), 340 m asl. It can only be reached by flying (or using a rope or a ladder for humans). The entrance room was filled with a quite undisturbed, 2 m thick sedimentary deposit with very rich small vertebrate bone assemblages deposited from barn owl pellets all along the sedimentation process.

These layers have been sampled extensively using special techniques to avoid contamination between the very dense upper layers and the lower ones [43,44]. Stratigraphical analysis and accelerator radiocarbon dating on the bones themselves both pointed to a rapid sedimentation rate during the last 2500 years, with a break in the building up of sediment between 1960 and 610 cal BP. The record is thus concerning two periods, from the Iron Age to the Late Classical Antiquity on one hand, and from the Central Middle Ages to the present times, with a gap for the Early Middle Ages, between the 4th and the 11th c. CE. Excavations and collection of the microvertebrate remains were conducted by J.-D. Vigne from 1989 to 1992 according to a very strict protocol [43,44].

Multivariate analyses of the frequencies of the small mammal species show that modern species had approximately the same ecology throughout the time sequence, and give, for the first time, an idea of the ecological preferences of the extinct endemic species of Corsica [45]. The general development of the faunal spectra shows a general increase in human impact (clearance of the vegetation), through the sediment sequence. Other multivariate analyses indicate a sequence of at least three agricultural cycles: the first one during the Early Roman period, the second one between the Late Roman period (2–5th centuries CE) and the Pisa Peace (11–13th centuries CE), and the last one during the 14–19th centuries CE. The two earlier ones were mainly concerned with cereal cultivation, the later with tree cultivation [43,45,46]. Whilst most of these cycles have only a local significance, two phenomena have a more extensive significance: (1) the immigration of *Rattus rattus* to Corsica and probably to the whole North-Western Mediterranean Basin, dated between the 4th–2nd centuries cal-bc; and (2) the mass extinction of the small endemic species, dating either to the Late Roman period or to the 11–13th centuries CE and probably resulting from large scale agricultural deforestation [47]. In addition, the small mammal assemblages recovered appear to reflect a wetter climate at both the beginning of the sediment sequence (the "small alpine glacial age") and at the end (14–18th centuries: "*Petit Âge Glaciaire*"[44]).

This site gives an example of the importance of small mammal assemblages for reconstructing the environment in the historical period and of well-documented stratigraphic accumulations for precisely dating the arrival of an invasive species such as the black rat.

Six samples were screened for aDNA analysis, all from 15th C deposits. Of these, two were selected for deeper sequencing (MDT001, MDT002).

***Petronell - Carnuntum Zivilstadt, Austria*** *(Günther Karl Kunst & Silvia Radbauer)*

The civilian town of Carnuntum (today's municipality of Petronell-Carnuntum) was located directly on the Pannonian *limes*. It developed since the 1st century CE, about 2.2 km to the west of the military camp, and became the administrative centre of the Roman province *Pannonia superior* from the beginning of the 2nd century CE onwards (municipium Aelium / colonia Septima). Important urban developments started in the Hadrianic and Severan periods and lasted until the 5th century CE [48].

In the southeastern part of the civilian town, a typical insula (so-called *insula VI*), delimited by four stone-paved streets, has been excavated and presented by the Archaeological Park Carnuntum (about 1.4 ha) for over seventy years. In 2002-2003, extensive archaeological excavations were carried out in most of the preserved sections of the western street (so-called *Weststraße*) of *insula VI* (comprising the baths and a so-called villa urbana). The excavations revealed a multiphase construction of the street (phases 1-4) with different stages of the associated aquatic infrastructure. The system became established by the end of the 1st/beginning of the 2nd century CE and remained in use until the final abandonment of the freshwater and sewage systems in the last decades of the 3rd century [49,50]. Both the layers of the street body and the backfilled or clogged features of the aquatic infrastructure produced rich bioarchaeological samples and various kinds of artefacts. According to recent excavations in *insula VI*, the use of the stone-paved surface of the street lasted well into the late Roman era (Phase 5).

About 40 remains of black rat were collected from the various substructures of the so-called *Weststraße* and from features pertaining to the adjacent buildings to the west, together with other animal bones and archaeological artefacts. Altogether four skull fragments and four mandibles, seven innominate bones and various long bones could be identified as belonging to *Rattus*. It is likely that some of these bones derive from the same skeletons. Eleven black rat femora representing discrete individuals were screened for aDNA within this study, from the contexts listed below. Of these, three samples were selected for deeper sequencing. Both specimens with fused and unfused proximal and distal epiphyses were included. The stratigraphic origin of these specimens can be described as follows.

A total of 16 rat bones, including four right femora, were found in the backfill layers of the fresh water channel K46 (RM 31-36, 52-56), which are mostly related to the abandonment of this system, when overlaying sediments from the Hadrianic and Severan periods broke in from above after the wooden beddings had collapsed. Thus, the material from the backfill may include a long timespan reaching from the middle of the 2nd to the final abandonment of the water supply system during the last decades of the 3rd century CE (phase 4) *(FN 9003/2003, SE 904: **DNA 125-229**; FN 9011/2003, SE 904; FN 9014/2003, SE 910; FN 10008/2003, SE 1004)*. Sample PZA002 derived from FN 9003/2003 and was directly dated to 125-229 CE (95.4%; MAMS-46371). Sample PZA003 derived from FN 10008/2003.

Unfortunately, the only right femur from the main sewer K14 (RM 62-74) originates from sediments relocated during earlier excavations *(FN 3010/2002, SE 325: **DNA 129-235**)*. Sample PZA001 derived from this context and was directly dated to 129-235 CE (95.4%; Wk-51523). The slight discrepancy between dates may relate to a small reservoir effect ($\delta^{13}C$ = -18.03). It should be mentioned, however, that the intact layers of the main sewer K14 (RM 42-61), both the lower sandy fill and the mudflow above, produced several rat remains including almost complete skulls. This sedimentary sequence developed as soon as the sewer was no longer maintained towards the end of the 3$^{rd}$ century CE [50].

In fact, most of the rat bones from the *Weststraße*, including those taken for analysis, derive from layers of phase 4, associated with the abandonment of the sanitary system occurring during the last decades of the 3$^{rd}$ century.

In some parts of the adjacent buildings to the west (RM 9-14) – located nearby a street latrine – two right femora from layers of the 2$^{nd}$ and 3$^{rd}$ century CE were found *(FN 6019/2003, SE 617; FN 6045/2003, SE 636)*.

***Old probostry, Prague Castle, Prague, Czech Republic*** *(René Kyselý)*

The analysed rat bone finds originate from the third courtyard of Prague Castle, from the archaeological excavation in house No. 48 (also known as the Old probostry) excavated in 1984 by I. Boháčová, J. Frolík and J. Žegklitz of the Institute of Archaeology of the Czech Academy of Sciences [51–55]. The animal bone assemblage was analysed in the 1980s by J. Petříčková [53]. The excavated area was the residence of the bishop of Prague until at least the end of the 13th century, and after this it

served as the probostry until the 19th century. As such the site is considered to represent a high social class for much of its occupation. The nature of the site's occupation between the 14th and 15th centuries is unclear, however, and it is possible that during this period the house was abandoned.
Eleven rat bones were screened here, six found in trench S III, feature 1, layer 45 (sample ID 174), and five in trench S VI, part of a waste pit, layer 230 (sample ID 229). One bone from each context was selected for deep sequencing. Altogether 76 rat bones were recovered from these two contexts, together with other animal bones and archaeological material. Sample 174 is dated based on stratigraphy and associated finds to the 10th or first half of the 11th century CE until 1061, and sample 229 to the 14th–15th century CE. One of the screened rat bones from each context was radiocarbon dated, returning dates of 1321-1417 CE (sample 174) and 1277-1387 CE (sample 229) respectively at 95.4% (OxA-39167, 571±19; Wk-51524, 679±21). The result of the first radiocarbon analysis (OxA-39167) seems to re-date the earliest rat occurrence in the Old probostry originally given in previous publications [53]. The group of variously aged individuals labelled 174 probably represents later intrusion – black rats which got themselves into the older context.
Nonetheless, these finds represent the earliest Holocene occurrence of *Rattus* in the Czech Republic confirmed by radiocarbon analysis. Approximately 15 archaeological sites/excavations in Bohemia have provided rat bones so far. At present no find of *Rattus* in the country is archaeologically dated to before the 9th-10th century CE. A number of black rat mummies have been recovered in other parts of Prague Castle, and appear to have been trapped in walls between 1541 and 1572 [56].

***Panga ya Saidi, Kenya (Richard Helm)***

Panga ya Saidi is an extensive karstic cave complex located on the Dzitsoni Uplands of Kilifi County in coastal Kenya. The two rat specimens derive from two test pits excavated in 2010 as part of an initial scoping survey, Test pit 1 Context 105 Spit I and Test Pit 2 Context 206 Spit G [57]. Further excavations have revealed a record of human activity extending back ~78ka years BP [58]. The test pits were excavated in adjoining chambers with collapsed ceilings open to the surrounding environment. Both specimens were recovered from aceramic deposits containing human habitation waste (ash, charred plant remains, animal bone and shell) associated with LSA stone-tool technologies. The sample from test pit 1 (PYS 10/1 I) derived from a deposit equivalent to layer 9 in later excavations, from which radiocarbon (OxA-29431 and OxA-30146) and OSL (OSL-13) sampling returned a calibrated and Bayesian modelled age of ~33ka years BP.

***Rirha, Morocco*** *(Tarek Oueslati)*

The diachronic site of Rirha is located inland about 80Km eastward from the capital Rabat. The site is settled within a meander of wadi Beht and comprises a Mauretanian Tell, a Roman city of about 10 ha and a medieval occupation. Recent excavations have been undertaken on a yearly basis since 2005 [59]. The chronological sequence ranges from the Mauretanian period (5th century BCE–1st century CE) to Roman times (40 CE–3rd century CE) and after a hiatus of over four centuries, two medieval phases were recognised (Idrisid 9th–10th century then Merinid 12th–14th century). The Mauretanian Tell, with its mud brick houses, streets, kitchens and dump areas has been excavated extensively and benefitted from sieving which provides clear evidence of the presence of black rat and house mouse in 1st century BCE levels and probably earlier around the 3rd 2nd c BCE for house mouse [60]. It is worth stressing that most black rat bones were hand collected during excavation as they appeared as whole or incomplete skeletons probably killed by the occupants of the Tell or predators cat and weasel being documented in the bone assemblage.

***Santa Maria Chapel, Lavezzi, France*** *(Jean-Denis Vigne)*

The Lavezzi archipelago consists of six main islets plus many emerging granite blocks, offshore to the far south-east extremity of Corsica. The southernmost islet is Lavezzi (66 ha). It is also the second most remote one, located at 3.5 km from the coasts of Corsica and at the same distance from the central islet of the archipelago, Cavallo (112 ha).

Intensive pedestrian surveys, small soundings and one excavation conducted under the direction of J.-D. Vigne [61] together with the study of many historical records on the Lavezzi island since the Classical Antiquity, allowed documentation of the use of the islet by humans during the last 2000 years and their impact on the abundant but fragile fauna of birds and mammals that lived on it during that times [43,46,61]. Most of the archaeological and archaeozoological information came from the excavation of the fillings of the small abandoned roman chapel of Santa Maria Lavezzi, built sometimes between the 10th and the 12th century CE. This chapel was rapidly deconsecrated and successively then sporadically occupied by herders, fishers or pirates. They severely impacted the wildlife of the island, especially birds [62] via intensive predation. During the same time, the chapel was also frequented by barn owls that left abundant accumulations of small mammal and bird bones [61,63].

The main layers of the small stratigraphic accumulation date to the 14th and 17th century. They were sealed by a thick deposit from the collapse of the chapel roof in the early 18th century. Both provided a large series of mouse (*Mus m. domesticus*) and black rat (*R. rattus*) teeth and bones.

The mice were only represented in the 14th century layer; mice were absent from the 17th layer and have not been recorded on Lavezzi during the 20th century. This indicated that the 14th century mouse population was extirpated at some point between the 14th and 17th century CE without subsequent recolonisation; it seems that the level of anthropic frequentation of the islet since the 14th century was not intense enough to allow the presence of a permanent commensal population of mouse [63].

More than 2000 specimens of black rat were studied. They indicated that their remains were not only accumulated by the barn owl, but also by humans that cooked and consumed them, and by dogs (bones in some coprolites; Vigne et al. 1994). Morphometric analyses indicated no difference between the 14th and the 17th C accumulations, but also a significantly larger body size of both of them with reference to the modern populations of South Corsica and of the Lavezzi islet itself. Vigne et al. (1993) concluded that this 14-17th century population of black rat had probably evolved in isolation during several centuries, acquiring a large before becoming extinct at some point between the 18th and 20th century. The extant population colonized the islet after the 18th century and has not been on the islet long enough to display detectable insularity syndrome.

Nine rat specimens were screened for aDNA, all from the 14th century deposit. Of these, 5 were selected for deeper sequencing: SML001, SML002, SML003, SML004, SML005.

***Sulzbach Castle, Germany*** *(Kerstin Pasda)*

Castle of Sulzbach, Sulzbach-Rosenberg, Landkreis Amberg-Sulzbach, Upper Palatinate, Bavaria, Germany. Archaeological excavations were carried out parallel to the renovation of the upper and lower castle of Sulzbach. The dating of the castle started in the 8$^{th}$ century and reached into the 17$^{th}$ century.

Based on archaeological results the earliest settlement activities at the castle hill can be narrowed down to at least the 8$^{th}$ century CE. The results prove the presence of a family which was a member of a late Carolingian or Ottonian aristocracy of the 9$^{th}$ and 10$^{th}$ century. A sarcophagus found at the castle may be the grave of the count Ernst († 865) who was one of the eminent henchman of the emperor Ludwig der Deutsche. Between 938 and 1003, the castle of Sulzbach was probably the principal caste of the margrave of Schweinfurt and so was probably one of the most important places in this area.

Due to a revolt of the count Heinrich von Schweinfurt against emperor Heinrich II in 1003 the power changed [64]. Probably as a consequence the castle went into the possess of the family of the Sulzbach earls. The high political significance of the Sulzbachs is visible in their marriage policy especially in the 11$^{th}$ and 12$^{th}$ century. 1130 Gebhard II (ca. 1125-1188 CE) married Mathilde, the daughter of the Welf duke Heinrich IX of Bavaria. Gertrud of Sulzbach was married 1131/32 to Konrad III of the Staufers dynasty. Her sister Luitgard of Sulzbach married 1138/39 the powerful Duke Gottfried of Lower Lorraine. Another Mathide of Sulzbach was married to the margrave Engelbert of Istria. Bertha of Sulzbach, also one of the sisters was married 1146 with the Byzantinia emperor Manuel I Komnenos and became after his death empress Irene of East Rome. In the 14$^{th}$ century Sulzbach became main capital of emperor Karl IV of his later so called “new boheminan” territory [65]. End of the 14$^{th}$ century Karl IV lost his interest in Sulzbach and the castle came into the possess of local Bavarian earls.

The periods of the castle were categorized according to archaeological construction phases [66]. The periods, which are of interest for the DNA-project are periods I-III (I: 8th to early 9th century; II: 9th to early 10th century; III: 10th century). All bone material was hand collected. The complete material consisted of 35003 animal remains and 29 single human bones. Only 11 bones could be identified as rats. Five of these bones were used for the project. Four bones were from layers that could be dated to the 8th to 9th century (period I-II), one to a layer dated to late 9th (?)/10th century (period I-III).

***Songo Mnara, Tanzania*** *(Stephanie Wynne-Jones and Jeffrey Fleisher)*

The site of Songo Mnara is a 14th – 16th century town, located on a small island off the southern coast of Tanzania[67]. It is part of a cluster of urban sites in the Kilwa archipelago, including the famous site of Kilwa Kisiwani. The town itself includes dozens of grand coral-built houses, five mosques and hundreds of tombs. It is one of the finest examples of the Swahili 'stonetown' tradition of the second millennium AD and it dates to the peak of Indian Ocean connectivity and trade along this coast.

The rat bones sampled here were excavated as part of a major campaign at the site between 2009 and 2016, which sampled domestic contexts across many different spaces. They come from a multi-layered midden deposit in the back room of House 44, one of Songo Mnara's coral-built houses. This was one of the richest contexts across the site, with ample evidence for cooking and food preparation. At Songo Mnara the assemblage of bones is dominated by fish (51% of total by weight). Rat bones were numerous at Songo Mnara and in this midden (21 NISP). The overall tetrapod assemblage is dominated by caprines and chicken, although a wide range of taxa are present representing a mixed economy. The context is interpreted as being a domestic midden that gradually accumulated over the 15th-century occupation, with several moments of clearing and covering, and the rats are considered intrusive.

***Tróia, Portugal*** *(Mariana Nabais)*

The Roman site of Tróia is located in the southwestern coast of Portugal, in a sandy peninsula delimited by the River Sado's estuary on the east, and the Atlantic Ocean on the west[68]. Such an environmental setting is highly favourable for fish and salt exploitation up to this day, and the site is believed to represent the largest fish-salting production centre known to the Roman Empire during the 1st and 2nd centuries AD [69–71].

Within the several identified fish-salting workshops, Workshop 2 is the earliest building dated in Tróia, from the Tiberian period, *i.e.* AD 14-37. The workshop has vats along its four walls displayed around a rectangular inner patio, and connected to Workshop 1 by a corridor. In the second quarter of the 3rd century AD, Workshop 2 was reorganised into two smaller workshops (2A and 2B), reflecting an economic retraction that affected the overall production capacity in Tróia, as well as in the whole southwest of Roman *Hispania* and the northwest of Morocco [72]. Nonetheless, there are indications of a production recovery in the first half of the 3rd century AD, which might have happened at a slightly earlier moment in Workshop 2B than everywhere else on site. During the first half of the 4th century AD, Workshop 2A enlarged its production capacity due to the incorporation of four extra vats that were initially from Workshop 1, therefore triplicating its production capacity. This last phase of production lasted until the second quarter of the 5th century AD [69–71].

The rat bones analysed here were recovered from the abandonment levels of vats 1 and 7c of Workshop 2, which were excavated in 2007 under the direction of Dr. Inês Vaz Pinto. The faunal remains were collected along with *Samian ware* and amphorae typical of the late 4th and early 5th centuries.

***Vila Franca do Campo, São Miguel, Azores*** *(Edward Treasure)*

The Azores archipelago consists of a group of nine volcanic islands situated in the North Atlantic Ocean, c.1400km west of the Iberian Peninsula and more than 2500km east of Newfoundland in North America. According to historical sources, the islands are first thought to have been discovered in the early-mid 15th century, with exploration of Santa Maria and São Miguel in 1431-2 [73]. Official colonisation of the islands is dated to 1439 when they were claimed by Henry the Navigator for the crown of Portugal[74]. However, it is possible that the islands were known and colonised before the 15th century, with palaeoecological evidence obtained from lake cores suggesting human settlement in the 13th century, if not earlier [75,76]. Despite this, it is from the 15th century onwards that human settlement of the islands intensified, notably on São Miguel which was the largest and most widely settled island within the archipelago [75,77].

On October 22nd, 1522, an earthquake struck the island of São Miguel, causing a major landslide above Vila Franca do Campo (henceforth Vila Franca); this was the most important settlement within the Azores. In 2015 and 2016, a series of small-scale excavations were undertaken within the footprint of the current city of Vila Franca to identify archaeological deposits associated with this landslide [78]. This involved the excavation of 26 trenches and some of these contained archaeological deposits directly associated with the 1522 landslide. The landslide was a highly distinctive and easily identifiable deposit. Due to the erosional nature of the landslide, no ‘intact’ settlement horizon was identified beneath it, but instead only the fragmentary remains of occupation deposits pre-dating 1522 were identified.

During the course of the excavations, animal bone was recovered through hand recovery, dry-sieving (1cm mesh) and bulk-sampling for flotation (0.5mm mesh). Bulk-sample residues were sieved into fractions (>4mm, 2mm, 1mm). The >2mm residue fractions were 100% sorted and, at a minimum, a sub-sample of the 1mm fraction (≥12.5%) was sorted. Rat bones were primarily extracted from these sample residues, although larger bones were recovered by coarse sieving during excavation. The rat bones were identified by Louisa Gidney and Sheila Hamilton-Dyer.

Four rat bones from Trench 25, Context [25027] were screened for aDNA, of which one (SMI001) was selected for deeper sequencing. This context represents debris pushed at the front of the landslide downhill towards the sea, and is one of a series of deposits containing high densities of artefactual and environmental remains (e.g. charcoal/charred plant remains, fish bone, animal bone). Pottery recovered from the deposits is consistent with a 15th to early 16th century date and numerous coins were also recovered, eight of which pre-date 1522.

***Voorburg - Forum Hadriani, Netherlands*** *(Jørn Zeiler)*

The Roman town of Forum Hadriani, situated in the present city of Voorburg (close to The Hague), was inhabited between 120 and 270 CE. From the early 19th C onwards there have been several excavation campaigns [79,80]. During one of these, carried out in 2005 by BAAC bv, a water well (feature no. 1015) made from a wooden barrel was excavated. It contained several hundred animal remains. Although the faunal assemblage was partly the same as in the rest of the site, with remains of cattle, sheep, pig, chicken, goose and mussels, there was a series of mammal and bird species that were not found anywhere else at the site.

Apart from polecat (skull and mandible), two bones of house sparrow and nine bones of raven there were 194 remains of rodents. A hundred of these could be identified: black rat (73), water vole (1), field vole (1), wood mouse (23) and house mouse (2). The rat bones comprised both cranial and postcranial elements and represented at least five individuals.

Of four right rat humeri from this well screened for aDNA, two were selected for deeper sequencing. One of the latter was also radiocarbon dated to 120-230 CE at 95.4% (Wk-51522, 1877±20), corroborating the archaeological date.

***York - Coppergate, England, UK*** *(David Orton)*

The site of 16-22 Coppergate was excavated by York Archaeological Trust between 1976 and 1981. Covering c.1000m$^2$, the excavations are famous for uncovering well-preserved remains of wooden buildings dating to Anglo-Scandinavian Jorvik, although Roman and later medieval deposits were also present. The site was subject to unusually intensive flotation sampling that produced a large assemblage of microvertebrate remains [81]. Rats appear throughout the Anglo-Scandinavian sequences but are particularly ubiquitous in Phase 3 (mid-late 9$^{th}$ to early 10$^{th}$ C), where they represent the earliest confident post-Roman finds of rat in York [82]. Rats are again frequent in Phase 5Cr/5Cf (mid to late 11th C). Nine rat specimens were screened for aDNA, mostly from Phase 3. Dates in Table S7 are based on revised phasing for the relevant contexts [83].

***York - Tanner Row, England, UK*** *(Terry O'Connor)*

The General Accident Extension site, 24-30 Tanner Row, York (YAT code 1983-4.32) was located to the South-West of the River Ouse, within the presumed Roman colonia. Excavations by York Archaeological Trust consisted of five linked trenches into 7.5m of archaeological deposit, much of it waterlogged and anoxic. In Roman levels, timber structures of late 2nd to early 3rd century were sealed by refuse deposition onto which mid-3rd century stone structures were built, continuing in use into the 4th century. Medieval activity mostly of the late 12th and 13th century consisted of refuse disposal onto and into an area of waste ground. The majority of animal bones recovered from the site were from the Roman timber levels, where preservation was excellent and context integrity generally good, and from medieval pits, with more oxidised preservation and more evidence of redeposition [84]. Wet-sieving was carried out for all levels of the site, with samples sieved on a 1mm mesh, then dry-sieved on 2mm mesh, from which all bone fragments were recovered.

A total of eight rat bones were screened for aDNA, deriving from four contexts dating securely to the mid-2nd to early 3rd centuries. Of these, three were selected for deeper sequencing: TRU001 and TRU002 from context 2457 and TRU003 from context 4155.

**Modern site descriptions**

***Zembra, Tunisia***

Modern rats were collected from the islet of Zembra by Jean-Denis Vigne in the 1980s, and subsequently skeletonised for use as reference specimens at the National Museum of Natural History, Paris. Eight individuals were screened here, and three selected for deeper sequencing (ZMB001, ZMB002, ZMB003).

**Supplementary Notes 2: *De novo* assembly of *Rattus rattus* reference genome**

**Autosomal chromosome assembly**

For the *de novo* assembly of *R. rattus* reference genome, we caught a male black rat individual from California, USA, where an invasive black rat population was established in the early 18th century [85]. This rat is now cataloged at the Museum of Vertebrate Zoology, UC Berkeley, https://arctos.database.museum/guid/MVZ:Mamm:236302. Genomic DNA was extracted from the liver and one shotgun sequencing library was prepared using Illumina TruSeq DNA PCR-free kit. The shotgun library was sequenced on Illumina HiSeq 4000 platform to produce 900M 150 bp paired-end sequencing reads, and a *de novo* assembly was generated using Meraculous [86], with k-mer size of 55, minimum k-mer frequency cutoff of 9 and diploid mode.

Three Chicago libraries and three Dovetail Hi-C libraries were further prepared following published procedures [87,88], and sequenced on Illumina HiSeq 4000 platform. A total of 498M PE150 reads were produced from the three Chicago libraries, providing an estimated physical coverage of 55.5X, which is the average number of read pairs of 1-100kb spanning a certain position in the genome. The Hi-C libraries were sequenced for 512M PE150 reads and provided an estimated physical coverage (10-10,000 kb pairs) of 15,256X. The *de novo* genome assembly from Meraculous and Chicago sequencing reads were first used as input data for HiRise scaffolding pipeline [87], then the output assembly, together with Hi-C sequencing reads were used for a second round of HiRise scaffolding.

**Sex chromosome assembly**

To retrieve the assembly from both sex chromosomes, we first mapped the shotgun sequencing reads to *Rattus norvegicus* reference genome Rnor_6.0 using BWA [89], and extracted reads mapped to the X chromosome (NC_005120.4) and Y chromosome (NC_024475.1), respectively.

Then we used Meraculous 2.2.5.1 [86] to get the *de novo* assembled scaffolds for both sex chromosomes, with k-mer size set to 55, minimum size cutoff as 9 and diploid mode disabled. The average and standard deviation of insert size was set to 400 bp and 10 bp, with average read length as 150 bp and the approximate genome sizes were 120 Mb for X chromosome and 2 Mb for Y chromosome. After Meraculous assembly, we reconstructed 9984 scaffolds from X chromosome, with a total length of 110.0 Mb and N50 length of 18.9 kb, and 440 scaffolds from Y chromosome, with a total length of 2.0 Mb and N50 length of 8.9 kb (Supplementary Table 1).

**Supplementary Table 1.** Meraculous *de novo* assembly of the sex chromosomes

| Chromosome | N Scaffold | Total length | Min length | Max length | N50 | L50 |
|---|---|---|---|---|---|---|

| | | | | | | |
|---|---|---|---|---|---|---|
| ChrX | 9984 | 110.0 Mb | 1.0 kb | 228.2 kb | 18.9Kb | 1644 |
| ChrY | 440 | 2.0 Mb | 1.0 kb | 45.7 kb | 8.9Kb | 69 |

These *de novo* scaffolds were then further assembled with Chicago and Hi-C data using HiRise genome assembly pipeline from Dovetail (Supplementary Table 2). After combing the sex chromosomes and autosomes assembly, we got the genome assembly of *R. rattus* consisted of 6805 scaffolds, with a total length of 2.25 Gb.

**Supplementary Table 2**. Statistics of the HiRise genome assembly

| | **Total length** | **N scaffold** | **N50** | **L50** | **N90** | **L90** |
|---|---|---|---|---|---|---|
| Autosomes | 2137.8 Mb | 5604 | 145.8 Mb | 6 | 73.3 Mb | 15 |
| ChrX | 110.9 Mb | 781 | 68.0 Mb | 1 | 1.5 Mb | 5 |
| ChrY | 2.1 Mb | 420 | 9.8 kb | 50 | 1.8 kb | 260 |

**Identification of scpMSY regions**

The single-copied scaffolds in the MSY region (scpMSY) were identified following a published strategy [90], using ten male and ten female *R. rattus* individuals with average genomic coverage over 1X (Supplementary Table 11). The mean coverage on each scaffold of each individual was calculated using reads with mapping quality and base quality over 30, and normalized by the mean coverage on all the 420 Y-chromosome scaffolds. The average normalized mean coverage (ANMC) of each scaffold was calculated as the average of normalized coverage in ten male individuals. The scaffolds with the average of coverage/nuclear mean coverage < 0.01 on the female individuals, and 0.1 < ANMC < 1.5 were considered as scpMSY scaffolds (Supplementary Table 14). Finally, we identified 321 scpMSY scaffolds, with a total length of 1,925,316 bp.

**Assessment of the genome assembly**

The repetitive regions were identified using RepeatMasker 4.0.7 (http://repeatmasker.org) [91] using Repbase 20170127 and the query species set as rattus (Supplementary Table 3), and TRF 4.09 (Tandem repeats finder) [92], with parameters set as “2 7 7 80 10 50 12”.

**Supplementary Table 3**. RepeatMasker summary for the *R. rattus* genome assembly

| **Percentage of sequence (%)** |
|---|

| | |
|---|---|
| Total | 38.34 |
| SINEs | 7.21 |
| LINEs | 16.39 |
| LTR elements | 9.87 |
| DNA elements | 1.24 |
| Unclassified | 0.4 |
| Small RNA | 0.03 |
| Satellites | 0.02 |
| Simple repeats | 2.87 |
| Low complexity | 0.33 |

The completeness of genome assembly was assessed by BUSCO 3.0.2 [93], using the 303 orthologs in Eukaryota odb9 dataset, and compared to the *Rnor_6.0* reference genome (Supplementary Table 4).

**Supplementary Table 4**. Comparison of BUSCO output between *R. rattus* genome assembly and *Rnor_6.0*

| **Assembly** | | **Complete** | **Fragmented** | **Missing** | **Total** |
|---|---|---|---|---|---|
| *R. rattus* | unmask | 273 | 11 | 19 | 303 |
| | mask | 269 | 13 | 21 | 303 |
| *Rnor_6.0* | unmask | 277 | 10 | 16 | 303 |
| | mask | 270 | 14 | 19 | 303 |

We aligned the new genome assembly with Rnor_6.0 reference genome using nucmer 4.0.0 in MUMmer tool package [94], to investigate the synteny between *R. rattus* and *R. norvegicus* genomes, using both masked assemblies and anchor matches that are unique in both reference and query. The 18 longest *R. rattus* scaffolds in autosomal assembly and 20 *R. norvegicus* chromosomes were well aligned (Supplementary Figure 1). As described in previous karyotype study, *R. rattus* has a diploid number of 2n=38. Chr5/7 and chr9/11 of *R. norvegicus* correspond to chr1 and chr2 of *R. rattus*, respectively [95]. The 9 largest scaffolds (>0.5 Mb) of the X chromosome assembly were also aligned to *R. norvegicus* chrX (Supplementary Figure 2).

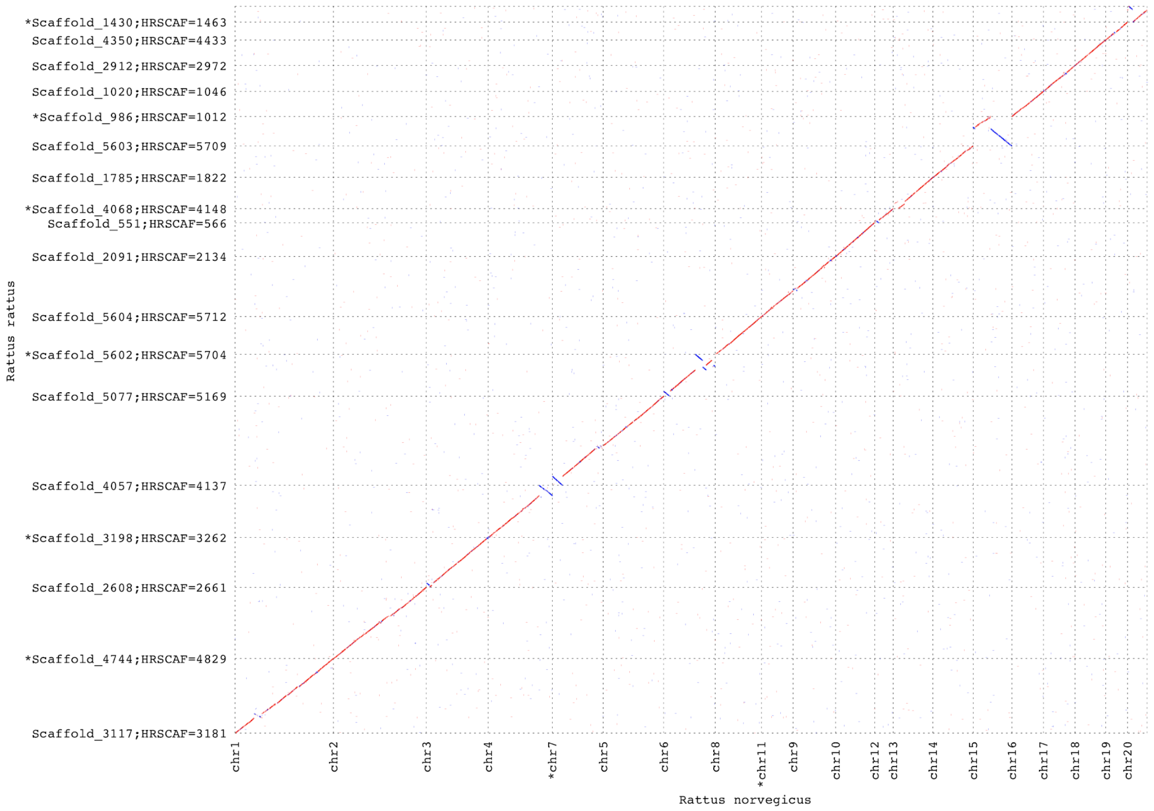


**Supplementary Figure 1.** Pairwise alignment between the *R. rattus* genome assembly and *Rnor_6.0* on autosomes.

The X-axis shows 20 autosomal chromosomes of Rnor_6.0 and the Y-axis shows the 18 largest autosomal scaffolds of the new genome assembly, each corresponding to one autosomal chromosome of the black rat. Forward matches are shown in red and reverse matches are shown in blue. The scaffolds with asterisk marked on Y-axis are plotted in a flipped orientation.

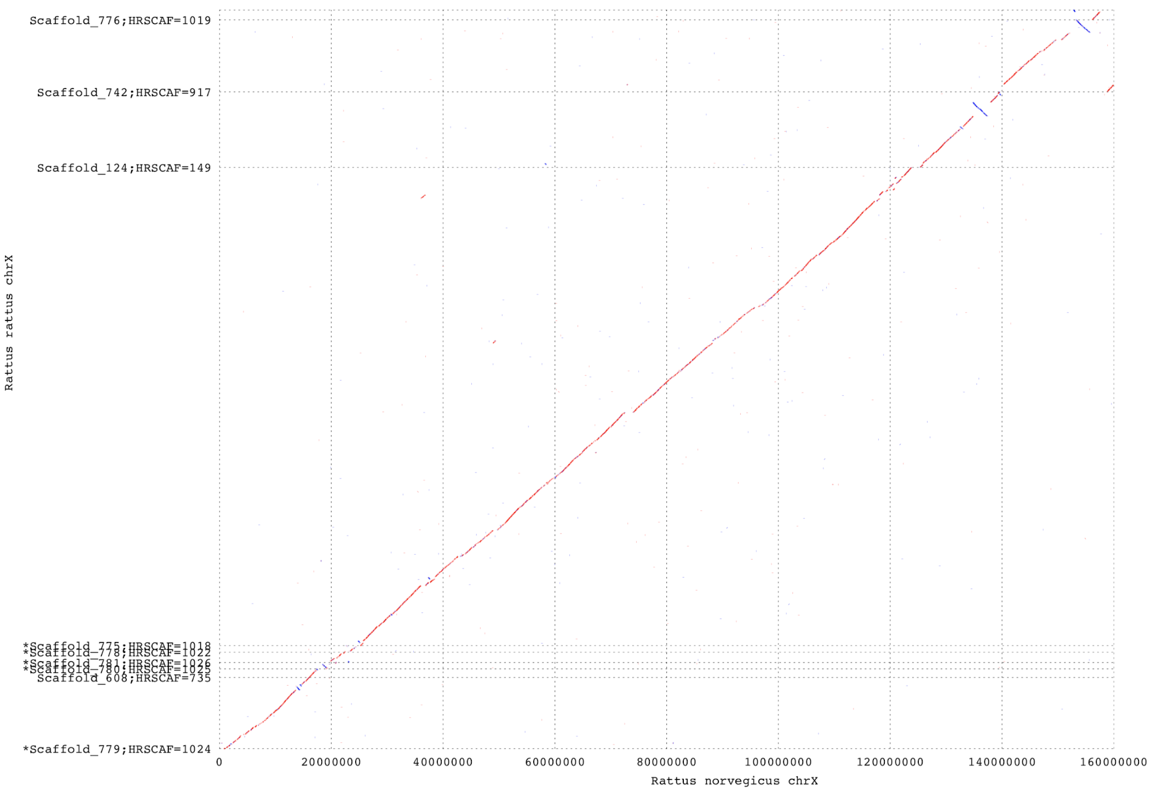


**Supplementary Figure 2.** Pairwise alignment between the *R. rattus* genome assembly and *Rnor_6.0* on chrX.

The X-axis shows X chromosome of *Rnor_6.0* and the Y-axis shows the 9 scaffolds over 0.5Mb of the new X-chromosome assembly. Forward matches are shown in red and reverse matches are shown in blue. The scaffolds with asterisk marked on Y-axis are plotted in a flipped orientation.

**Supplementary Note 3: Demographic analysis using G-PhoCS**

The Generalized Phylogenetic Coalescent Sampler (G-PhoCS) was applied to estimate the population sizes, population divergence times and migration rates among the three rat species [96], using high-coverage, diploid genomes. After masking out the repetitive regions identified by RepeatMasker and TRF, we identified 38,078 loci of 1kb length on autosomal regions, with less than 10% masked sites and interlocus distance over 50 kb, allowing for sufficient recombination.

The G-PhoCS analysis was performed based on a given topology (norvegicus, (rattus, tanezumi)), using the modern black rat CP-5999 and ERS215791, HXM4 from published study to represent for *R. rattus*, *R. norvegicus* and *R. tanezumi*, respectively. We first ran a preliminary analysis with all eight possible migration bands added in the model, for 250,000 generations and sampled every 100 generations. The prior for all migration events set to alpha=0.001, beta=0.0001 and all theta and tau parameters were set to alpha=3, beta=1000. The result was checked and summarized with Tracer 1.6.0 [97] and the first 50,000 generations were burn-in. In this preliminary run we identified only one migration event with total migration rate (m_total=m*tau) over 0.01, that is, the gene flow from rattus/tanezumi ancestral population into norvegicus lineage (Supplementary Table 5).

Based on the preliminary results, we performed two parallel runs for 500,000 generations and sampled every 100 generations, with one migration event and priors for theta and tau parameters set as Supplementary Table 6. Finally, the estimated parameters were converted to effective population sizes (Ne), divergence times (T) and total migration rates (m_total) as described in Gronau *et al.* [96]: theta = 4*Ne*μ, tau = T*μ/g and m_total=m*tau, with mutation rate $\mu=2.96*10^{-9}$ site/generation and generation time (g) of 0.5 years.

### Supplementary Note 4: Taxonomy of the black rat and mitochondrial phylogeny based on cytb region

#### Taxonomy of the black rat

*Rattus rattus* (also known as the black rat, roof rat or ship rat) and *Rattus tanezumi* belong to the '*Rattus rattus* species complex' within *Rattus*; these species and their relatives all form an extremely taxonomically complicated set of closely related (and interrelated, see below) taxa belonging to a number of genera (the Rattini tribe) within Rodentia. The *Rattus* genus comprises approximately 66 currently recognised species [98]. It is the most diverse genus of rodents and the largest of mammalian genera with a range of highly adaptive specialists and also multiple generalists (Rowe et al., 2011). The fossil and subfossil record for *Rattus* is sparse and an inability to identify between closely related species precludes confident assessment of ancestral taxa [99,100]. As a result, assessment of species divergences and radiations within *Rattus* using the fossil record has generally only been possible on the large scales, such as the assessment to the arrival of *Rattus* to New Guinea and Australia (e.g. see Sahulian *Rattus* [101]). Biomolecular and morphological advances have helped clarify the taxonomic complexity within this genus and its closest relatives [100–104], but questions still hang over the association of many of these species; for example *Bandicota* sp. and *Rattus* sp. [103].

Within the *Rattus* genus a number of lineages and/or species form the *Rattus rattus* species complex [103,105]. This is possibly the most complicated species complex within mammals, with more than 150 different species names having been previously assigned to populations and lineages within it, the resulting number of synonyms continue to cause taxonomic confusion in modern studies [106]. Biomolecular methods (when considered as encompassing both single marker studies and multi marker studies) have helped resolve a number of these taxonomic problems, but in a number of instances, where they have used single or few markers (e.g. studies using only mtDNA) they have revealed even greater complexity, and raised numerous questions, which directly reflects the nature of these species' evolutionary past. For example, the *Rattus* genus harbours a number of deeply divergent lineages, some reflect likely real taxonomic division (e.g. the split between *Rattus rattus* and *Rattus tanezumi* as examined here) but others reflect deeply divergent lineages that exist within single populations of the same species (e.g. the finding of the mtDNA lineage for *Rattus* clade 3 [103], which microsatellite evidence showed had no taxonomic support [107]). Even more complicated still are species within the *Rattus rattus* species complex that appear to have diverged more recently than most lineages within the complex, yet have developed distinct morphological differences and correspondingly show little to no signs of gene flow; therefore forming distinct taxonomic units. For example, *Rattus sakeratensis* (formerly one population of a number previously identified as *Rattus losea* [103,106]) appears to have rapidly diverged and is now considered a distinct species [107]. In light of the complex evolution of these taxa, resolving the systematics of *Rattus* requires in-depth biomolecular and genomic approaches.

Within the *Rattus rattus* species complex the two major commensal taxa with the most extensive human associated distributions are *Rattus rattus* (identified as mtDNA lineage I in [105], see overview below in section on mitochondrial phylogeny) and *Rattus tanezumi* (the east Asian house rat, identified as mtDNA lineage II in [105]). *Rattus rattus* (the primary subject of this study) is widely distributed and appears to be associated with global trade routes associated with European imperialism; *Rattus tanezumi* has an almost exclusively east Asian distribution, east of the Tibetan plateau [105]. These two species represent some taxonomic uncertainty, both appear morphologically similar, though with some variation in pelage colour [108]. However, pelage colour (coat colour of belly and/or back) is highly variable both among populations, but also within population; furthermore, taxonomic divisions based on coat colour within *Rattus rattus* have led to some of the proliferation of weak taxonomic divisions [98,103,105,106]. An initial basis for division, and most likely a restriction on gene-flow between these lineages stem from chromosomal rearrangements [109–111]. Chromosomal variation is often considered an unsatisfactory criteria for species definition in mammals, but can be considered an indicator of potential accelerated rates of speciation and a possible restriction of gene-flow (Ayala & Coluzzi, 2005; Corti & Rohlf, 2001; Navarro & Barton, 2003; Rieseberg, 2001; Saïd et al., 1999; Searle, 1998); within *Rattus* the majority of species are 2N=42, which is as a result thought to be the ancestral arrangement for the genus [112]. *Rattus tanezumi* has a karyotype arrangement of 2N=42; in contrast *Rattus rattus* (lineage I) has at least 3 karyotype races, but the primary configuration is 2N=38; an additional karyotype race in Mauritius has the same number as the ancestral form (2N=42), though in a different arrangement; and a further race with 2N=40 is found in Sri Lanka [109,110,113–117]. However, these differences in karyotype arrangement are not a complete gene-flow barrier and in fact the two mtDNA lineages associated with each taxon, *Rattus rattus* and *Rattus tanezumi*, are found in anthropogenically introduced and mixed populations in Sri Lanka, South Africa, California and Japan [105,118–120]. With evidence for admixture between karyotype races and animals with the separate mtDNA showing free gene-flow in human introduced populations, the question of the taxonomic division between *Rattus rattus* and *Rattus tanezumi* requires a greater and in-depth analysis of multiple lines of evidence to test this division; here we examine whole genomic evidence.

**Earliest *Rattus rattus* commensalism and its western natural occurrence: Natufian rats**

The presence of *Rattus sp.* (cf. *Rattus rattus*) is often cited as an indication of a shift to more sedentary settlements of the Natufian period (~15,000–10,000BP) in the Levant region [121,122]. If correct, these *Rattus* specimens would be the furthest and earliest western range of *Rattus* post the Last Glacial Maximum (~20,000±2,000 BP for Europe, prior to 20,000–40,000BP a rat species cf. *Rattus haasi* is reported from central and western Eurasia, but its relationship to other major *Rattus* species is unclear [121]). Furthermore, the link between Natufian *Rattus rattus* with Natufian sedentism is based upon the suggestion that these early *Rattus rattus* finds are also commensal, which would also make these the earliest examples of commensal *Rattus*. In this way *Rattus rattus* finds are combined with a suite of other likely commensal species including house mouse (*Mus musculus*), and house sparrows (*Passer domesticus*) to bolster support for Natufian shifts towards sedentary settlements and the first steps towards domestication of cereal grains [123]. However, amongst these species displaying early commensal behaviour, *Rattus rattus* remains are usually recovered in the smallest numbers and most sporadically and none of these *Rattus rattus* remains have been directly dated [122]. Therefore, although this might represent some of the earliest steps of *Rattus rattus* toward commensal behaviour the earliest evidence for intense and large scale commensal behaviour appears to be from Indus Valley sites [99].

**Mitochondrial phylogeny based on cytb region**

For an overview of the relationship between the ancient rats and modern black rats from across their range, 292 tissue samples of *R. rattus* were collected and analysed for mitochondrial cytb region. Among them 263 specimens were obtained from various museums (Field Museum, Chicago; American Museum of Natural History, New York; British National History Museum, London) and additional 29 modern specimens were collected in the field by Dr. J. Chris Hillman. The museum material is composed mainly of dried tissue, skins or ethanol-fixed samples, modern specimens were stored in ethanol. The sampling area comprises different places of the mainland and islands around the Indian Ocean, including countries from the East and West African coast, the Arabian Peninsula, as well as the Indian subcontinent and South-East Asia.

DNA extraction was performed in dedicated ancient and modern DNA laboratories in the Archaeology Department at Durham University. The different sample types (wet and dried tissue, skin) used in this analysis were prepared prior to DNA extraction in order to minimise the risk of coextracting exogenous contaminants and to remove preservative chemicals that can inhibit subsequent PCR, respectively. Dried skin samples were washed several times with Tween 20, ethanol- or formalin-fixed specimens were washed several times with purified water in order to increase the quality of DNA recovery. Afterwards, samples were placed on a petri dish and finely chopped with a sterile, disposable blade before transferral of ~ 10 mg into a 1.5 ml Eppendorf tube. The different

DNA extraction protocols for each sample type are stated below. One in ten extractions were blank controls in order to detect possible contamination.

Extraction protocol 1: Dried tissue

DNA extraction of dried tissue samples was carried out using the Qiagen MicroKit, following the manufacturer's recommendations.

Extraction protocol 2: Ethanol-fixed tissue

DNA extraction of ethanol-fixed tissue samples was carried out using the following protocol.

Day 1

Add 300 µl Extraction buffer (1M NaCl, Tris-HCl pH 8.0, 10% SDS, H2 O) to each sample, add 3 µl Proteinase K to each sample, vortex 15 sec, incubate samples on a rotary shaker overnight at 50°C

Day 2

Add 80 µl of saturated NaCl to remove DNA byproducts, vortex centrifuge 10 min at 9.000 rpm, transfer supernatant without touching the pellet, discard pellet repeat step until the supernatant is clear.

Add 800 µl ethanol (97-100%) to precipitate the DNA pellet, mix by inverting the tubes several times, centrifuge 45 min at 13.000 rpm.

Pour off the supernatant of each sample and remove any fluids with a small pipette, but mind the DNA pellet. Leave the tube open to dry the pellet, wait until all fluids are dissolved.

Add 200 µl 1xTE, incubate 5 min, centrifuge 1 min at 14.000 rpm, end up with one elution E1 á 200 µl, freeze at -20°C until further use.

PCR protocol

The cytochrome B region of the mitochondrial DNA was targeted for PCR amplification. PCR set-up was conducted under a fume hood in a pre-PCR clean room. Every PCR set-up included a negative control in order to detect possible contamination. Additionally, a positive control (modern material of each particular species) was included in each PCR to exclude that possible failure of the reaction is due to reagents or the thermal cycler. The modern positive control was stored in the dedicated post-PCR room and added to the reaction before placing the tubes in the thermal cycler.

Standard protocols of PCR set-up and thermal cycler programs are stated below. To ensure optimal PCR success, modification of the reaction conditions was repeatedly needed. Modification included altering the amount of DNA extract added or adjusting the final concentration of the other reagents used. Furthermore, cycling conditions were changed by in- or decreasing the times and numbers of cycle repetitions. PCRs were visualised on a 1.5% agarose gel, using GelRed and UVillumination. Successfully amplified PCR products were stored at -20°C prior to sequencing.

The first set of primer pairs U1/L1 to U4/L4 has been designed by Trinks. The second set of primer pairs Cyt b Rr1 to Cyt b Rr10 have been designed in equal parts by Eager and Trinks. Primer pairs F1 and F2 were taken from Aplin et al. (2011). The design is based on the sequence of a whole mitochondrial genome of *Rattus rattus* (NCBI accession number NC_012374).

**Supplementary Table 19**. Primers for mitochondrial fragment amplification.

| Fragment Ref_NC_012374 | Primer forward 5'-3' | Primer Position | | Primer reverse 5'-3' | Primer Position | | Product Length | °C |
|---|---|---|---|---|---|---|---|---|
| Cyt b U1/L1 | AATTTGTCATTATTTCTACACAGCATT | 14043 | 14069 | TAGGGTTGCTTTGTCTACTGAGAA | 14628 | 14652 | 559 bp | 56 |
| Cyt b U2/L2 | CATCTGCCGAGACGTAAACTAC | 14330 | 14352 | GTCTCCTAGTAAGTCTGGGAAGAAT | 14858 | 14883 | 507 bp | 56 |
| Cyt b U3/L3 | AGGATCAAACAACCCCACAG | 14735 | 14755 | TGTTGATGGTGGGGAGTTAGT | 15353 | 15374 | 599 bp | 56 |
| Aplin Museum F1 (2011) | ATCACACCCTCTACTCAAAA | 14144 | 14164 | GGCATGTAAGTATCGRATTAG | 14358 | 14378 | 194 bp | 56 |
| Aplin Museum F2 (2011) | TCATCAGTTACYCACATCTGC | 14316 | 14337 | CCTCAGATTCATTCGACTAGRGT | 14601 | 14624 | 264 bp | 56 |
| Cytb b Rr1 | ACACAGCATTTAACTGTGACCA | 14060 | 14082 | GGGCGGGAAGGTCAATGAAGG | 14176 | 14197 | 94 bp | 56 |
| Cytb b Rr2 | TTAATCACTCCTTCATTGACCTTCC | 14167 | 14192 | AGCCGTAGTTTACGTCTCGGCAG | 14333 | 14356 | 141 bp | 56 |
| Cytb b Rr3 | TTAACAGCATTCTCATCAGTTAC | 14304 | 14327 | GTTGCTATGACTGCAAATA | 14485 | 14504 | 158 bp | 56 |
| Cytb b Rr4 | TCCTACACCTTCTTAGAAACATGAAAC | 14442 | 14469 | AGCCTCCTCAGATTCATTCGAC | 14607 | 14629 | 138 bp | 56 |
| Cytb b Rr5 | CAAACCTATTATCAGCCATTCCCTA | 14566 | 14590 | AGTTTAGTCCTGTGGGGTTGTT | 14742 | 14764 | 151 bp | 56 |
| Cytb b Rr6 | GCCCTTGCAATTGTACATCTCCT | 14697 | 14720 | TGGGTCTCCTAGTAAGTCTGGGAA | 14862 | 14886 | 142 bp | 56 |
| Cytb b Rr7 | GACTTACTTGGAGTATTCATGTTAC | 14808 | 14833 | GGGATGGAGCGTAGAATAGCG | 14960 | 14980 | 127 bp | 56 |
| Cytb b Rr8 | ACCCCACCACATATTAAGCCAGA | 14916 | 14939 | TGGGCGGAATGTTAGACTGCGT | 15062 | 15084 | 123 bp | 56 |
| Cytb b Rr9 | TTCTAATCTTAGCCTTTCTACCA | 15019 | 15042 | AACTRATGGATGCTAGTTGG | 15179 | 15199 | 137 bp | 56 |
| Cytb b Rr10 | AGCCAACCTCTTCATTTTAAC | 15113 | 15134 | GCTCTTCATTTTTGGTTTACAA | 15300 | 15322 | 166 bp | 56 |

All samples were analysed in the facilities at the Archaeology Department of Durham University. Because of the recent age of most of the specimens, the samples were treated as modern material. Each workstep – from DNA extraction to sequencing set-up – was conducted in the modern laboratory, following standard extraction protocols. The sequencing reaction was carried out by the DNA Sequencing Service at the School of Biological and Biomedical Sciences at Durham University. Mitochondrial DNA was amplified in 10 overlapping fragments for cytochrome b, whereas a variety of primer combinations was used depending on the nature of the sample. The sequencing chromatograms were edited manually, subsequently assembled, and a consensus sequence per individual exported using Geneious R6 version 6.0.6 (Drummond et al. 2011). Standard anti-contamination guidelines were followed.

The phylogenetic maximum likelihood tree of the cytb region revealed all the ancient rats from this study belong to the black rat lineage (Supplementary Figure 4). We observed the same substructure as Aplin with five black rat haplogroups. In addition to these five haplogroups we have discovered a new group which consists of modern samples from Sri Lanka and the Andaman Islands and is basal to all black rats in this study. We have named these haplogroups A through F.

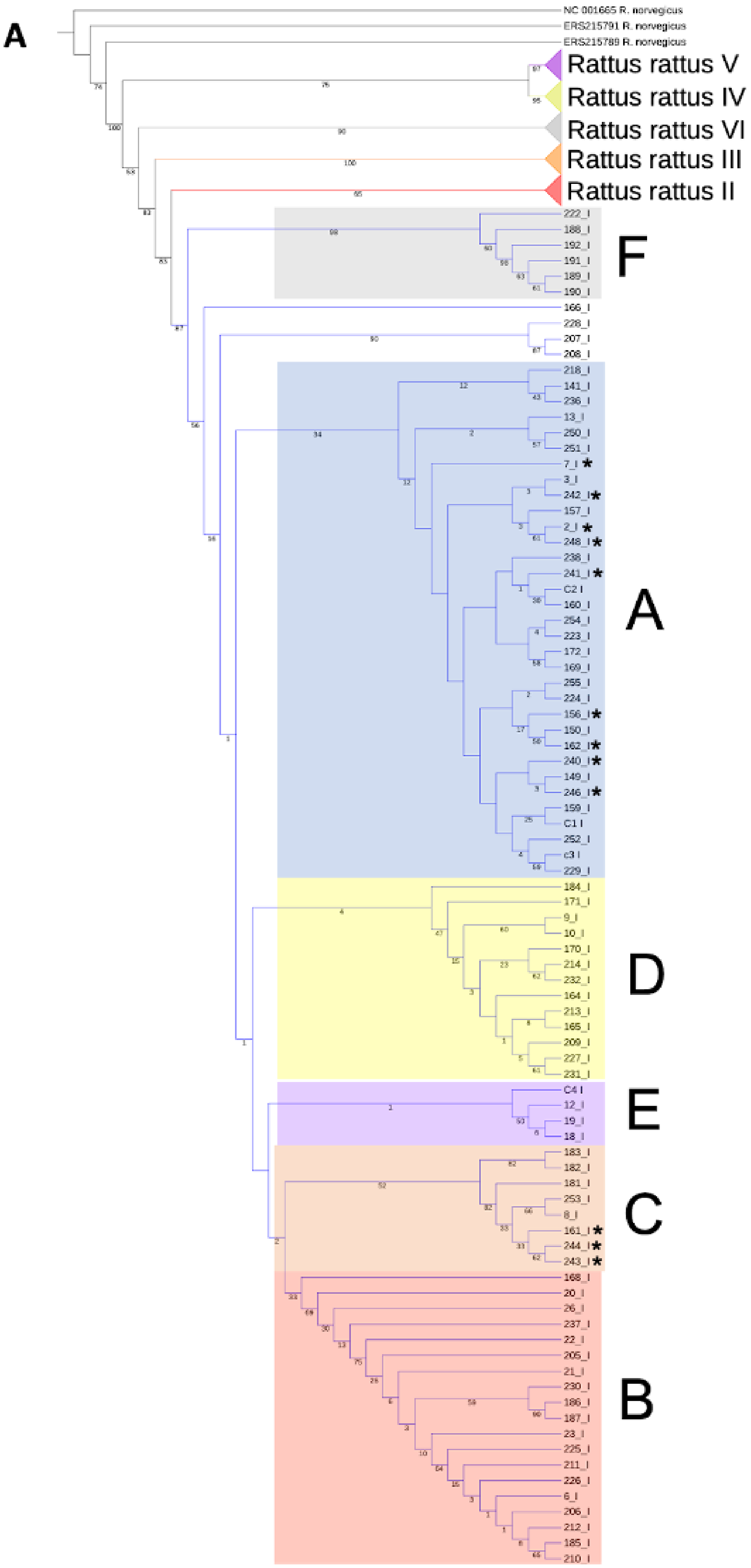
A
NC 001665 R. norvegicus
ERS215791 R. norvegicus
ERS215789 R. norvegicus
Rattus rattus V
Rattus rattus IV
Rattus rattus VI
Rattus rattus III
Rattus rattus II
F
A
D
E
C
B

**B**

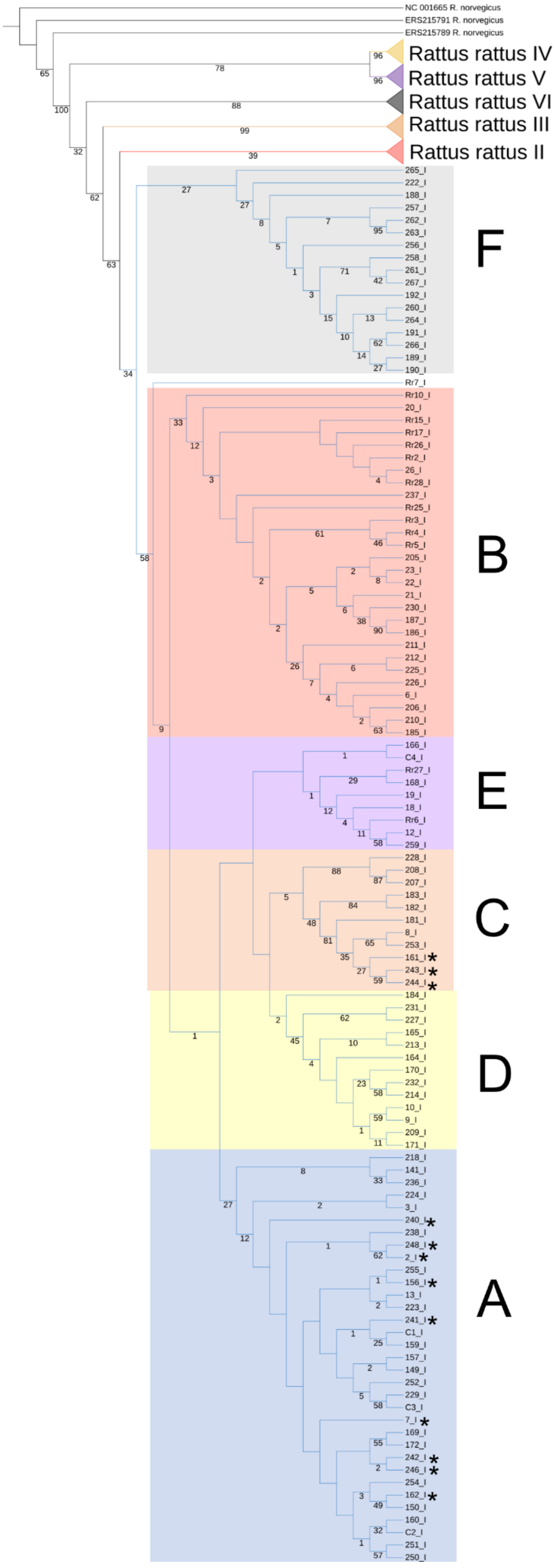
NC 001665 R. norvegicus
ERS215791 R. norvegicus
ERS215789 R. norvegicus
Rattus rattus IV
Rattus rattus V
Rattus rattus VI
Rattus rattus III
Rattus rattus II
265_I
222_I
188_I
257_I
262_I
263_I
256_I
258_I
261_I
267_I
192_I
260_I
264_I
191_I
266_I
189_I
190_I
F
Rr7_I
Rr10_I
20_I
Rr15_I
Rr17_I
Rr26_I
Rr2_I
26_I
Rr28_I
237_I
Rr25_I
Rr3_I
Rr4_I
Rr5_I
205_I
23_I
22_I
21_I
230_I
187_I
186_I
211_I
212_I
225_I
226_I
6_I
206_I
210_I
185_I
B
166_I
C4_I
Rr27_I
168_I
19_I
18_I
Rr6_I
12_I
259_I
E
228_I
208_I
207_I
183_I
182_I
181_I
8_I
253_I
161_I *
243_I *
244_I *
C
184_I
231_I
227_I
165_I
213_I
164_I
170_I
232_I
214_I
10_I
9_I
209_I
171_I
D
218_I
141_I
236_I
224_I
3_I
240_I *
238_I
248_I *
2_I *
255_I
156_I *
13_I
223_I
241_I *
C1_I
159_I
157_I
149_I
252_I
229_I
C3_I
7_I *
169_I
172_I
242_I *
246_I *
254_I
162_I *
150_I
160_I
C2_I
251_I
250_I
A

**Supplementary Figure 4.** Maximum likelihood tree (GTR+GAMMA) of rat samples based on CYTB region

A. This figure includes ancient samples from this study alongside haplotypes from Aplin, Colangelo, Etougbétché as well as modern samples from Trinks and Eager. See supplementary tables for a list of samples corresponding to each haplotype (411 sequences, 192 haplotypes in total). Haplotypes denoted with a * have at least one ancient sample found with this haplotype.

B. This figure includes samples above and the addition of modern samples from India (481 sequences, 214 haplotypes in total). Those from mainland India were not included in the main tree due to their sequence lengths being less than 90% however due to their importance we built this second tree. Haplotypes denoted with a * have at least one ancient sample found with this haplotype.

The map shows the six *Rattus rattus* complex groups and that the samples from our study reflect the same regional partitioning observed in Aplin et. al. [105]. Lineage I and lineage II are the only samples to be seen outside their natural range with lineage I being the main lineage moved around the world (Supplementary Figure 5).

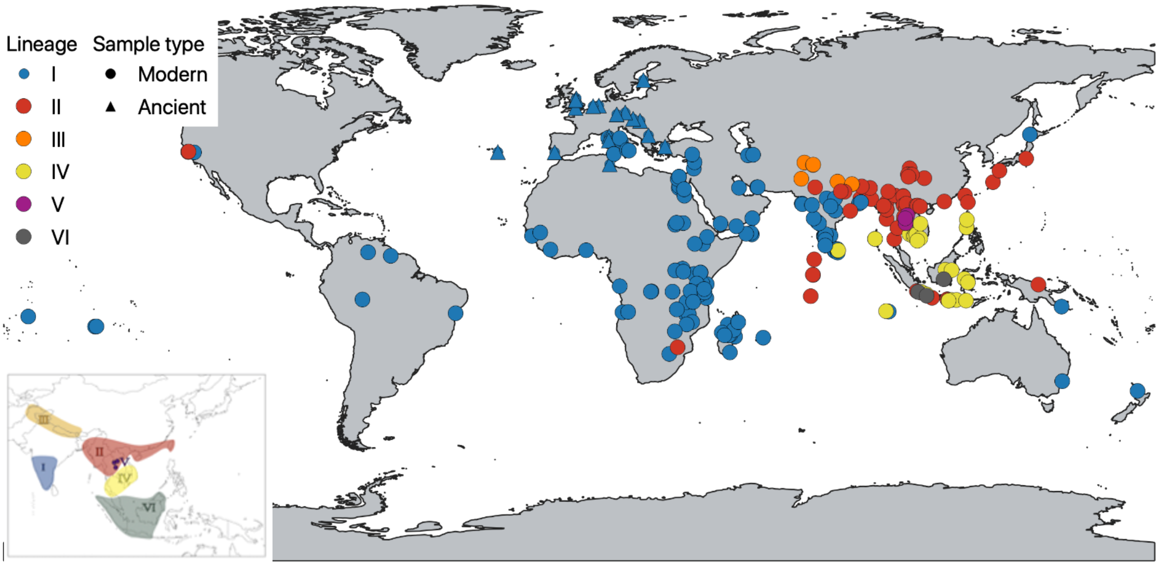


**Supplementary Figure 5.** Map showing the mitochondrial lineage included in the cytochrome B analysis.

Triangles depict ancient samples and circles modern, colours depict the different mitochondrial lineages within the *Rattus rattus* complex. The insert in the bottom left is an image from [105] showing the modern distribution of the *Rattus rattus* complex.

The median joining network largely reflects the same structure as the maximum likelihood tree with groupings of haplotypes by regions with a large star-like haplogroup containing haplogroup A (Supplementary Figure 6). All the haplotypes which are in haplogroup A are connected to the centre of the star by one or two substitutions. Any haplogroups differing by more than this are found on the maximum likelihood tree as a separate haplogroups. There are two other clear star-like clusters, one containing mostly samples from Madagascar (Hap 164) and the other largely from India and East Africa (Hap 6). The haplogroup containing Hap 189 is the most derived. The other grouping which is also derived is the haplogroup containing Hap 6.

**Supplementary Note 5: Whole genome analysis on ancient black rats**

**Estimation of heterozygosity**

To further investigate the genetic diversity of black rats through time and space, we estimated the heterozygosity of ancient rat genomes based on pseudo-diploid genotypes on the nuclear genome. Then we summarized the distributions across different time periods and regions (Supplementary Figure 7, Table 13). We found that in both Roman/Byzantine and medieval/post-medieval time periods, the rats from the southern group possessed higher heterozygosity than rats from the northern group. The low genetic diversity in the northern group could be explained by the founder effect of limited waves of introduction into this region, likely related to the Roman expansion [124–126], which also corresponded to the clustering of all northern group rats together in the phylogenetic analysis (Figure 3A). Conversely, the Mediterranean region has a longer history of rats, dating at least to the first millennium BCE [127,128]. Frequent contacts across the Mediterranean, and perhaps beyond to Asian regions with established rat populations, may have enabled multiple waves of introduction into different sampling areas, as revealed by the multiple lineages of southern group rats in the phylogenetic analysis (Figure 3A).

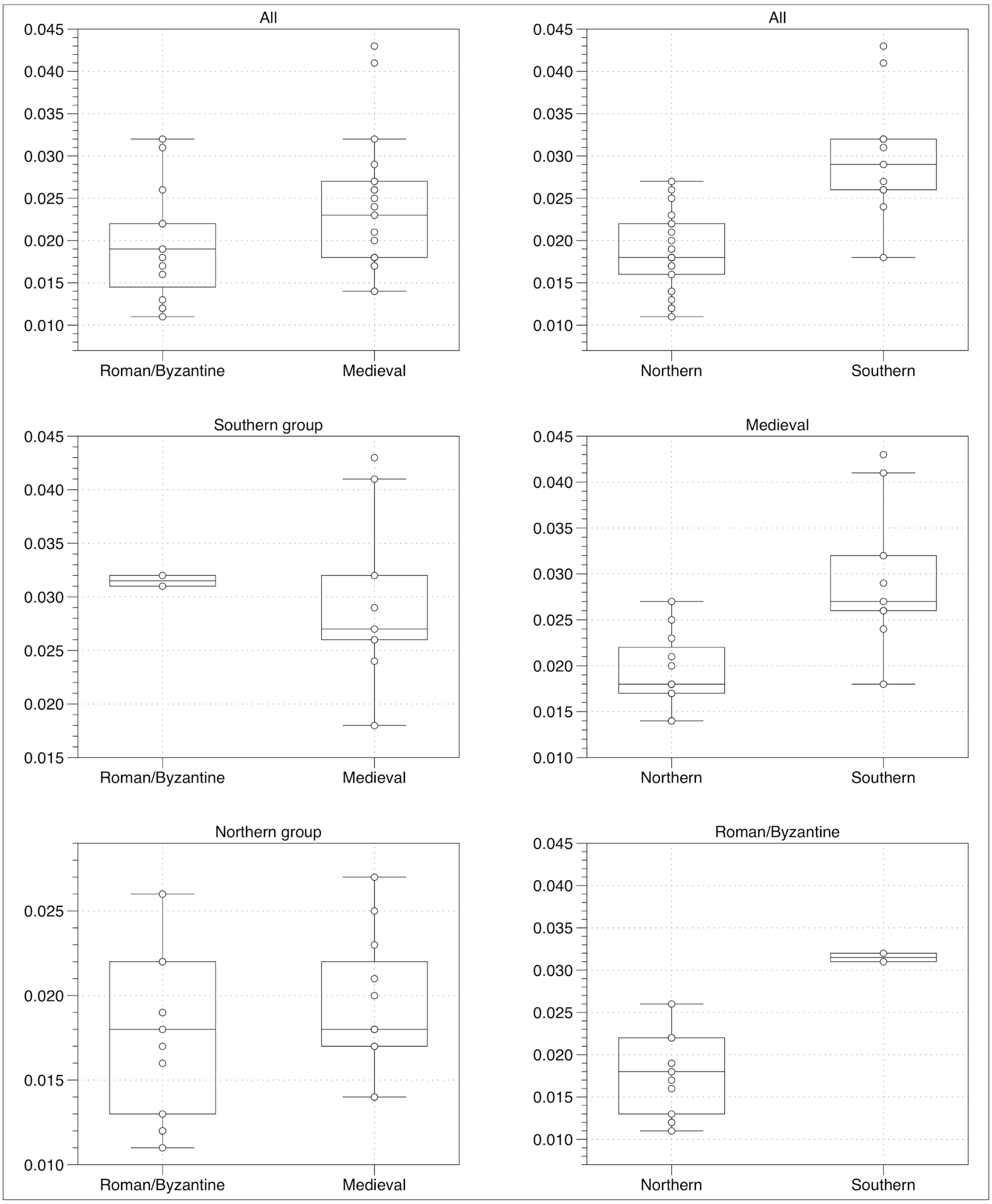


**Supplementary Figure 7.** Heterozygosity of ancient black rats on autosomal variants.
This figure summarizes the heterozygosity of each ancient individual estimated from pseudo-diploid genotype calling, grouped based on time periods and geographic regions.

**MDS plot based on nuclear genome variants**

The multidimensional scaling (MDS) plot based on isolation-by-state (IBS) distance among these samples also revealed a similar pattern. When the outgroups were included, the black rat, Asian house rat and brown rat samples were clearly distinguished by the first two dimensions (Supplementary Figure 8). In the MDS plot consisting only of the black rat samples, the first two dimensions separated three Mediterranean island groups (Zembra, Lavezzi archipelago and Corsica) from the major cluster, while the third and fourth dimensions revealed the distinction between Roman/Byzantine and medieval/post-medieval samples from temperate Europe.

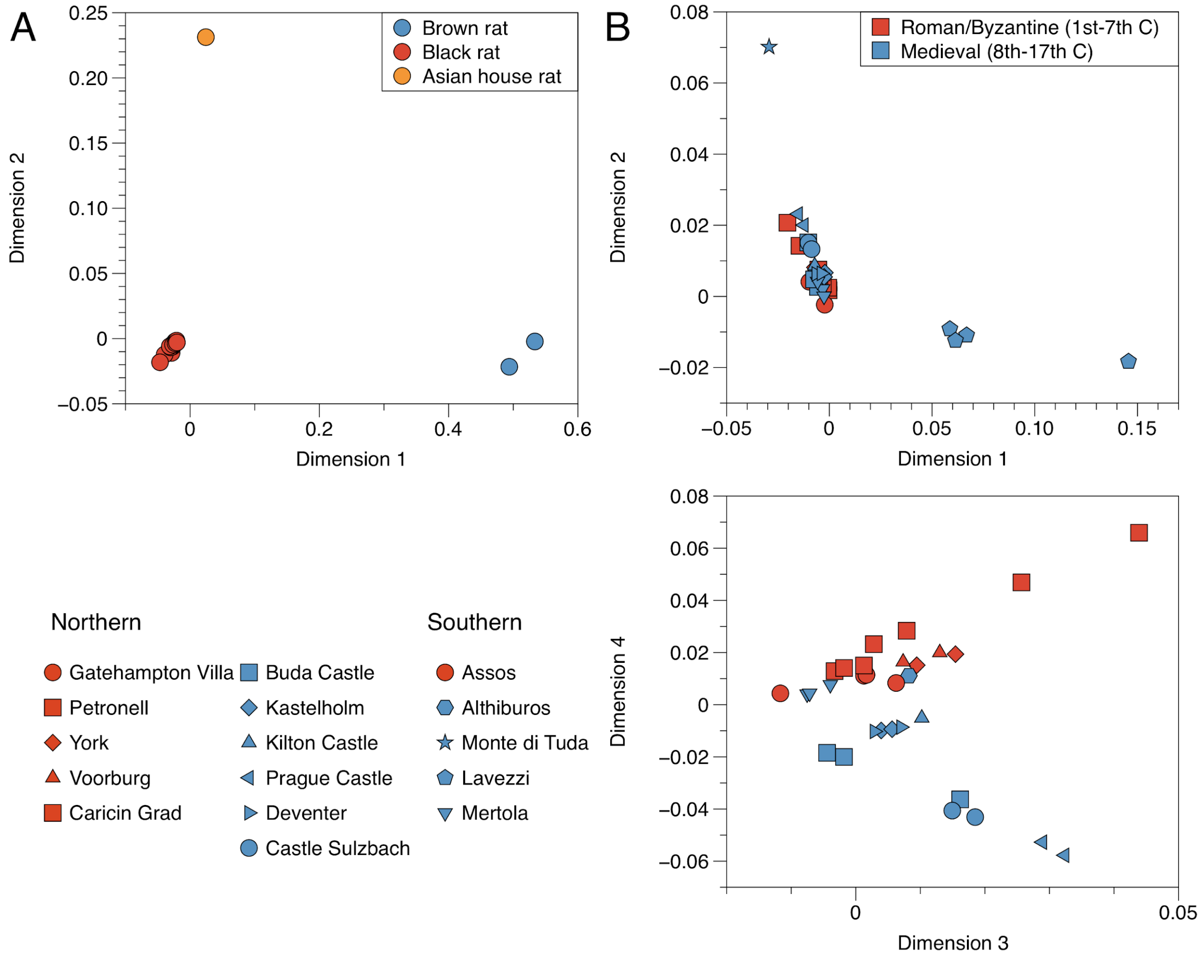


**Supplementary Figure 8.** MDS plot of the rat samples based on IBS distances.
**(A)** The plot of all rat samples including modern and ancient black rats (red), modern brown rats (red) and the Asian house rat (orange). The first two dimensions clearly distinguish the three rat species.
**(B)** The plot of ancient black rats. The first two dimensions distinguish the southern populations from the others and the third and fourth dimensions further show the distinction between Roman/Byzantine and medieval populations from the northern group.

#### Assessment of the *R. tanezumi* introgression in the modern black rat

The modern *R. rattus* sample (CP-5999) sequenced for reference genome assembly was collected from California, USA, where admixture between *R. rattus* and *R. tanezumi* was proposed by microsatellite studies [118,119]. To test whether the CP-5999 sample carried introgression from *R.*

*tanezumi*, we applied $f_4$-statistics in the form of $f_4$(norvegicus, tanezumi; CP-5999, Testpop) for all the ancient rat groups (Supplementary Table 19). We found that the modern rat is equally related to *R. tanezumi* with all ancient rats, ruling out the possibility of genetic introgression in this individual.

**Supplementary Table 20**. $F_4$-statistics of the affinity to *R. tanezumi* in black rats

| Testpop | F4 | Z | nBABA | nABBA | nSNPs |
|---|---|---|---|---|---|
| Ass | -0.000011 | -0.283 | 27364 | 27442 | 6835117 |
| ATU | -0.000011 | -0.236 | 12710 | 12754 | 4072243 |
| BUD0014 | 0.000044 | 1.151 | 19356 | 19110 | 5592353 |
| BUD003 | 0.000053 | 1.154 | 12420 | 12218 | 3810370 |
| Car | -0.000044 | -1.12 | 28332 | 28641 | 6973691 |
| GAU | -0.000009 | -0.203 | 8852 | 8882 | 3244499 |
| KAF | -0.000038 | -0.878 | 12086 | 12233 | 3920611 |
| KLT | -0.000031 | -0.683 | 14471 | 14607 | 4326576 |
| MDT | -0.000092 | -2.129 | 16759 | 17200 | 4783967 |
| MEP | -0.000025 | -0.692 | 18856 | 18997 | 5679673 |
| PRA | -0.000026 | -0.632 | 17674 | 17811 | 5221123 |
| PZA | -0.000057 | -1.428 | 14156 | 14419 | 4592312 |
| SML | -0.000091 | -2.26 | 27357 | 27980 | 6819026 |
| SNE | -0.000045 | -0.792 | 10224 | 10366 | 3120163 |
| Sulz | -0.000025 | -0.6 | 23211 | 23365 | 6225331 |
| TRU | -0.000047 | -1.107 | 16714 | 16958 | 5157166 |
| VOB | -0.000011 | -0.261 | 15900 | 15955 | 5081887 |
| ZMB | -0.000089 | -2.129 | 26124 | 26707 | 6593161 |

### Supplementary figures

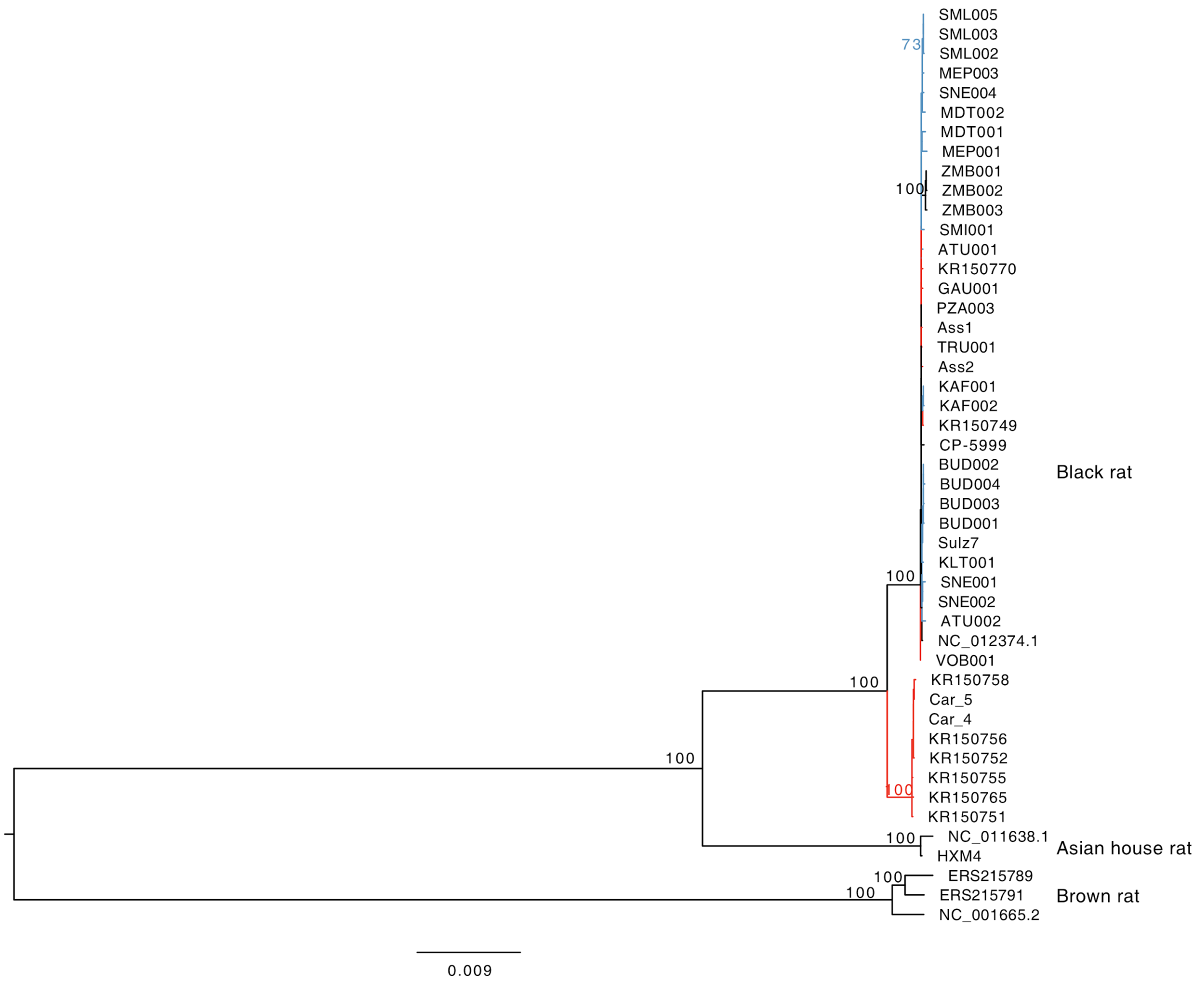


**Supplementary Figure 3.** Phylogenetic relationship among ancient black rats on mitochondrial genomes.

The maximum likelihood tree is constructed using the brown rat as outgroup, with bootstrap support from 100 replicates marked on each node. The species names corresponding to three clusters are written on the right side. The branches with red or blue colors correspond to ancient black rats from Roman/Byzantine and medieval periods.

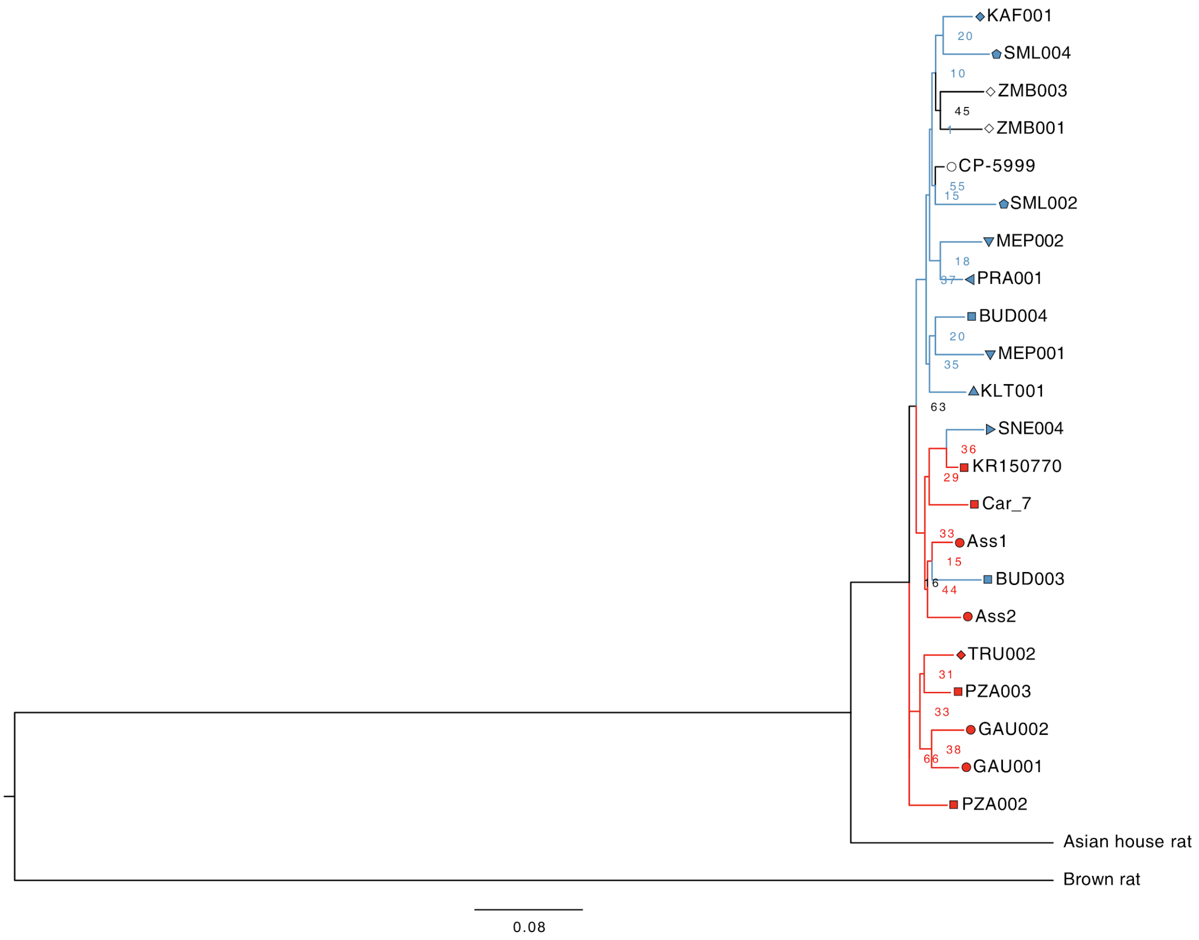


**Supplementary Figure 9.** Phylogenetic relationship among ancient black rats on Y-chromosome. The maximum likelihood tree is constructed using the brown rat as outgroup, with bootstrap support from 100 replicates marked on each node. Except for one brown rat and one Asian house rat, all the black rats form a cluster. The cluster is further divided into two clades, one exclusively formed by Roman rats (red) and the other includes Byzantine (red), medieval (blue) and modern (black) rats.

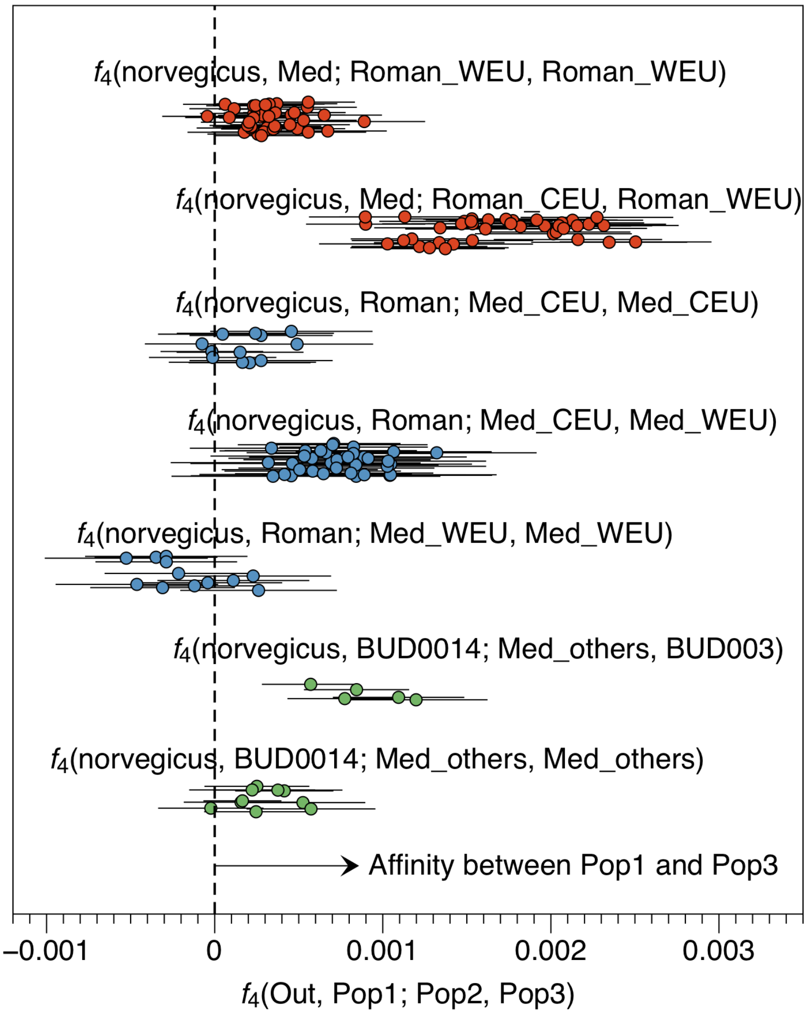


**Supplementary Figure 10.** The *f*-statistics showing admixture between different ancient rat populations

The dots show all the combinations of $f_4$-values as described above each cluster, and the error bars show ± 3SE of the estimates. Here the red clusters show that the medieval rats (Med) are closer related to western European Roman rats (Roman_WEU), compared to central European Roman rats. The blue clusters show that the Roman rats (Roman) are closer related to western European medieval rats (Med_WEU), compared to central European medieval rats (Med_CEU). The green clusters show that the post-medieval Buda Castle rats (BUD001/4) are closer related to the medieval Buda Castle rat (BUD003), compared to other medieval rats from continental Europe (Med_others).

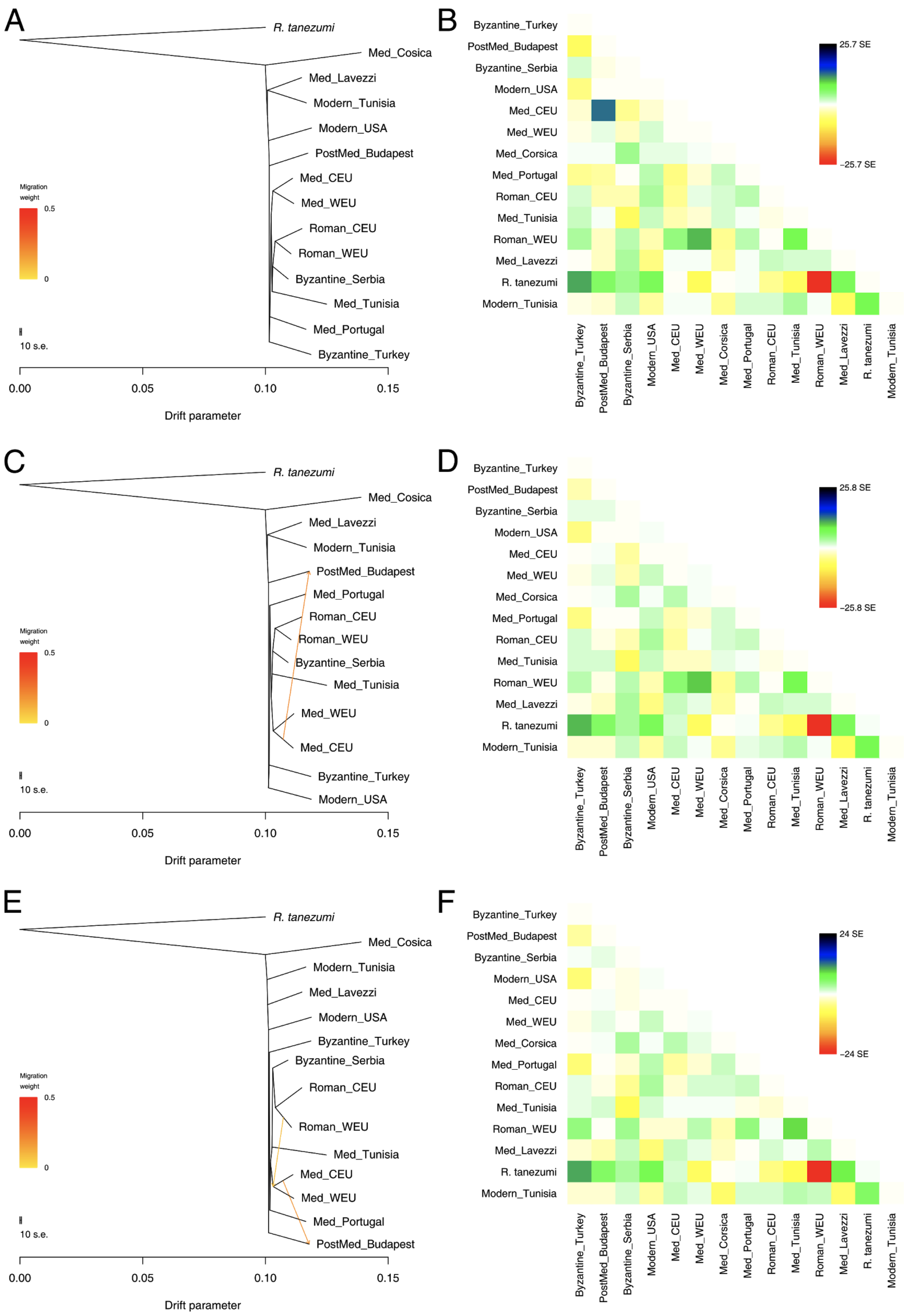


**Supplementary Figure 11.** Admixture graph among ancient black rat populations.

The ML trees estimated by Treemix with (A) no (C) one and (E) two migration edges fitted are plotted with the Asian house rat individual as outgroup. The corresponding residue covariance matrices are shown in (B), (D) and (F).
